# Microglial homeostasis requires balanced CSF-1/CSF-2 receptor signaling

**DOI:** 10.1101/2019.12.23.885186

**Authors:** Violeta Chitu, Fabrizio Biundo, Gabriel G. L. Shlager, Eun S. Park, Ping Wang, Maria E. Gulinello, Solen Gokhan, Harmony C. Ketchum, Kusumika Saha, Michael A. DeTure, Dennis W. Dickson, Zbignew K. Wszolek, Deyou Zheng, Andrew L. Croxford, Burkhard Becher, Daqian Sun, Mark F. Mehler, E. Richard Stanley

**Author notes:** Correspondence: Dr. E. Richard Stanley. ESP: MSB 7.147 Department of Neurosurgery, University of Texas Health Science Center at Houston, 6431 Fannin street, Houston, TX, 77030 USA. KS: Institut Cochin, INSERM U1016, CNRS UMR8104, Université Paris Descartes, Sorbonne Paris Cité, 27 rue du Faubourg St-Jacques, 75014, Paris, France.

## Abstract

*CSF-1R* haploinsufficiency causes adult-onset leukoencephalopathy with axonal spheroids and pigmented glia (ALSP). Previous studies in the *Csf1r^+/-^* mouse model of ALSP hypothesized a central role of elevated cerebral *Csf2* expression. Here we show that monoallelic deletion of *Csf2* rescues most behavioral deficits and histopathological changes in *Csf1r^+/-^* mice by preventing microgliosis and eliminating most microglial transcriptomic alterations, including those indicative of oxidative stress and demyelination. We also show elevation of *Csf2* transcripts and of several CSF-2 downstream targets in the brains of ALSP patients, demonstrating that the mechanisms identified in the mouse model are functional in man. Our data provide new insights into the mechanisms underlying ALSP. Since both increased *CSF2* levels and decreased microglial *Csf1r* expression have also been reported in Alzheimer’s disease and multiple sclerosis, we suggest that the unbalanced CSF-1R/CSF-2 signaling we describe in the present study may contribute to the pathogenesis of other neurodegenerative conditions.

**Highlights:**

- ALSP is a *CSF1R*-deficiency dementia associated with increased *CSF2* expression
- In *Csf1r+/-* ALSP mice CSF-2 promotes microgliosis by direct signaling in microglia
- Targeting *Csf2* improves cognition, myelination and normalizes microglial function
- CSF-2 is a therapeutic target in ALSP

## Introduction

The colony stimulating factor-1 receptor (CSF-1R) is regulated by two cognate ligands, CSF-1 and interleukin-34, is expressed on microglia and is required for their development and maintenance (reviewed in (Chitu et al., 2016)). Recent studies show that microglia play important roles in the regulation of neuronal development, learning-dependent synaptic pruning and oligodendrogenesis (Chitu et al., 2016, Hagemeyer et al., 2017). Thus, the microglial CSF-1R may non-cell autonomously regulate neural lineage cells.

Adult-onset leukoencephalopathy with axonal spheroids and pigmented glia (ALSP) also known as hereditary diffuse leukoencephalopathy with axonal spheroids and pigmented glia, pigmented orthochromatic leukodystrophy is an autosomal dominant, neurodegenerative disorder caused by mutations of the *CSF1R* gene (reviewed in (Konno et al., 2018)). ALSP is characterized by dementia with neuropsychiatric and motor deficits and has an average disease duration of 6.8 years. The discovery of an ALSP patient with a *CSF1R* frame-shift mutation that abolished CSF-1R protein expression proved that *CSF1R* haploinsufficiency is sufficient to cause ALSP (Konno et al., 2014).

Similar to ALSP patients, *Csf1r^+/-^* mice exhibit behavioral, radiologic, histopathologic and ultrastructural alterations associated with microgliosis and demyelination (Chitu et al., 2015). The microgliosis occurred in the absence of compensatory increase in the expression of CSF-1R ligands. However, in both presymptomatic and diseased mice, microgliosis was associated with an increase in the expression of mRNA of the proinflammatory microglial mitogen, colony stimulating factor-2 (CSF-2), also known as granulocyte macrophage CSF (GM-CSF) (Chitu et al., 2015). In the present study, we show that CSF2 expression is also increased in the brains of ALSP patients and we examine the effect of removal of a single *Csf2* allele on the development of ALSP in *Csf1r^+/-^* mice.

## Results

### Monoallelic Targeting of *Csf2* Prevents White Matter Microgliosis in Young *Csf1r^+/-^* Mice

As CSF-2 is a microglial mitogen (Lee et al., 1994) and *Csf1r^+/-^* mice exhibit early-onset microgliosis (Chitu et al., 2015), we initially examined the effect of genetic targeting of *Csf2* on Iba-1^+^ microglia density in young *Csf1r^+/-^* mice. Mono-allelic targeting was sufficient to normalize Csf2 mRNA expression and to lower the microglial density in *Csf1r^+/-^* mice to wild type levels (Fig. 1A-C). *Csf2* homozygous deletion showed no additional effects over *Csf2* heterozygosity. Consistent with the low abundance of *Csf2* transcripts in wild type brains (Fig 1C), Iba-1^+^ cell densities in *Csf2^+/-^* and *Csf2^-/-^* mice were not different from those of wild type mice (Fig. 1 A, B).

**Figure 1.**
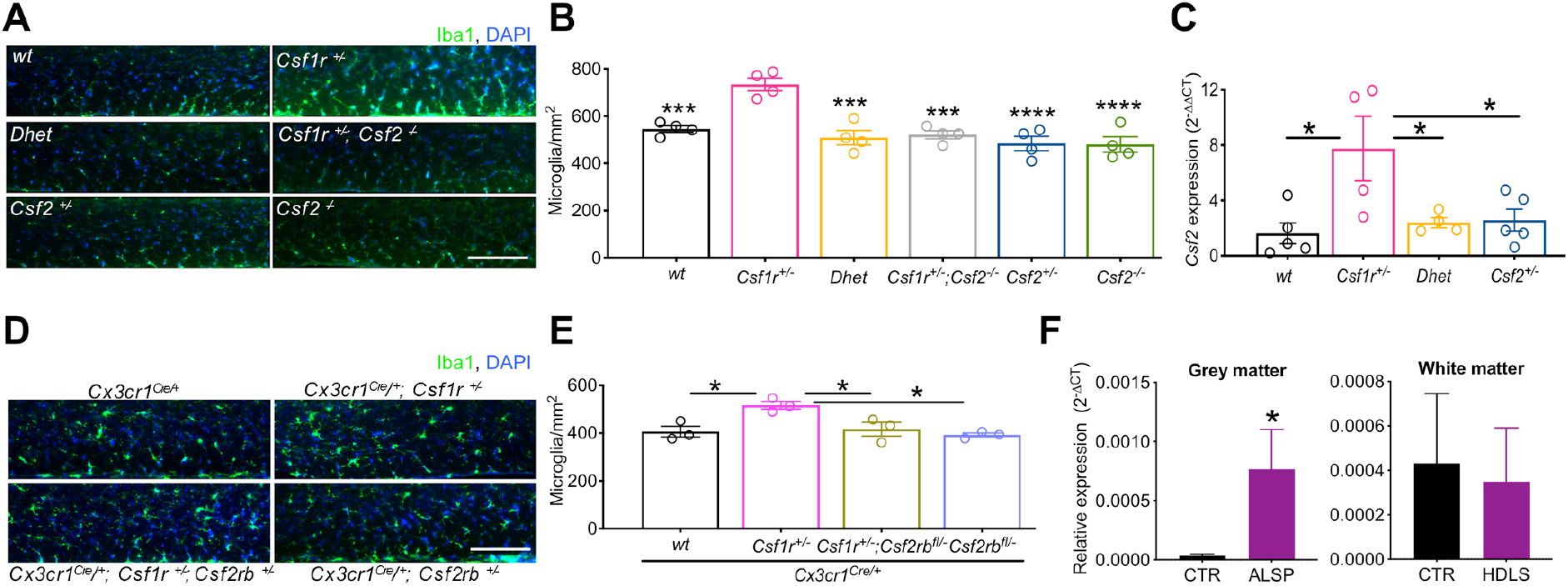
Involvement of CSF-2 in the Pathology of ALSP. (A) Microglial densities in the corpus callosum of 4-5-month-old mice. (B) Quantification of microglial densities. Two-way ANOVA followed by the Dunnett’s post hoc test. (C) Restoration of normal *Csf2* expression in 3-4-month-old *Csf1r^+/-^* mice by monallelic targeting of *Csf2*. One-way ANOVA followed by Tukey’s multiple comparison test. (D, E) Evidence for direct regulation of the increase of Iba1+ cell density by Csf2. Panel D, microglial densities; panel E, quantification. Data ± SEM, 3-month-old mice. One-way ANOVA followed by Tukey’s multiple comparison test. Scale bars, 100μm. (F) Expression of *CSF2* in the periventricular white matter and adjacent grey matter of ALSP patients and healthy controls determined by real time qPCR and normalized to *RPLl3*. N=5; *, p < 0.05, one-tailed Student’s t test. Data here and in all subsequent figures are presented as means ±SEM and only the significantly different changes are asterisked. See also Supplemental Figure S1 and Table 1.

To investigate whether CSF-2 drives microgliosis in ALSP mice directly, by stimulating its receptor in microglia, we used the *Cx3Cr1^Cre/+^* gene targeting system (Yona et al., 2013, Goldmann et al., 2013) to delete a *Csf2rb* allele (Croxford et al., 2015) in myeloid cells and microglia. The mice were examined at 3 months of age, when there is no evidence of demyelination or other pathological changes (Supplemental Fig. S1), that could confound the interpretation of the results. Similar to *Csf2* heterozygosity, conditional deletion of a single *Csf2rb* allele was sufficient to prevent white matter microgliosis in young *Csf1r^+/-^* mice without altering microglial densities in the wild type background (Fig.1D, E). These data indicate that CSF-2 causes microgliosis in the ALSP model via direct stimulation of microglia.

### Increased Expression of *CSF2* in the Brains of ALSP Patients

To investigate whether *CSF2* was elevated in the human disease, we examined the expression of *CSF2* in the periventricular white matter and adjacent grey matter of 5 ALSP patients and 5 matched controls (Supplemental Table 1). Consistent with the results obtained in mice, the levels of *CSF2* transcripts were almost undetectable in control patients, whereas *CSF2* expression was increased in the gray matter of the ALSP patients (Fig. 1F).

### Monoallelic *Csf2* Inactivation Prevents Loss of Spatial Memory in ALSP Mice

The normalization of microglial density in young mice prompted us to determine the effect of monoallelic *Csf2* inactivation on the development of behavioral deficits in older *Csf1r^+/-^* mice. Regardless of genotype, body weights of both males and females increased with age without significant genotype-associated differences (Supplemental Fig. S2) and there were no significant differences in survival up to 18 months of age. To test the effect of *Csf2* heterozygosity on short-term spatial memory, we used object recognition, object placement and Y-maze paradigms. In the object recognition test, deficits observed in *Csf1r* haploinsufficient mice at 7 months of age were prevented by *Csf2* heterozygosity (Dhet) (Fig. 2A). In the Y-maze (Fig. 2B) and object placement (Fig. 2C) tests, deficits apparent at 13-15 months in *Csf1r* haploinsufficient mice were also prevented by *Csf2* heterozygosity. Interestingly, in the latter two tests, deficits were also present in *Csf2* heterozygous mice. Similar results were obtained, at 16.5 months of age, in a test of long-term memory (object recognition with a 24h retention interval, Fig. 2D). Again, in these older mice, a deficit became apparent in the *Csf2* heterozygous mice. These results demonstrate prevention of short- and long-term spatial memory deficits of *Csf1r^+/-^* ALSP mice by *Csf2* heterozygosity. However, they also show that, *Csf2* heterozygosity on a wt background results in spatial memory deficits with aging.

**Figure 2.**
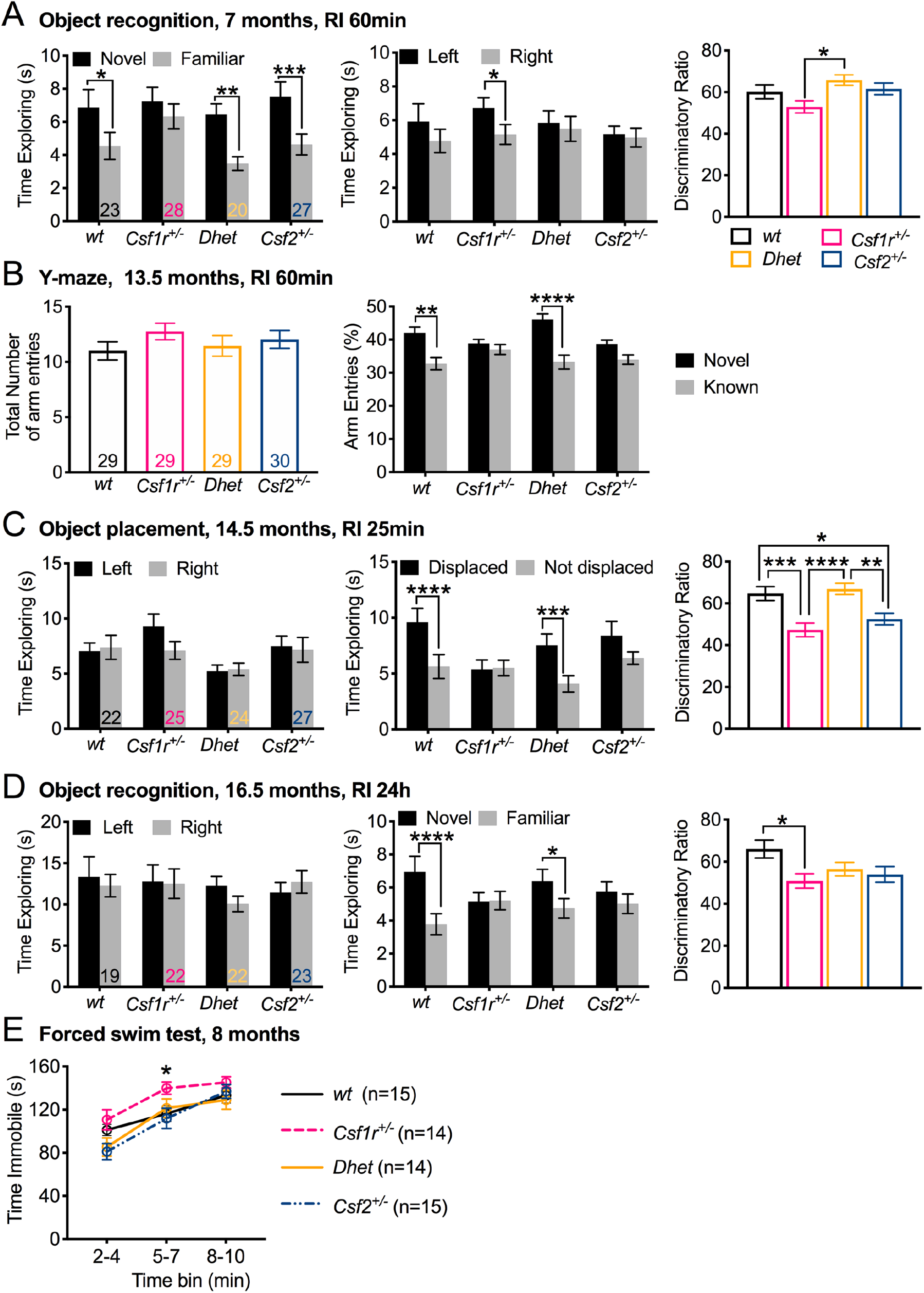
Deletion of a Single *Csf2* Allele Prevents the Cognitive Deficit and Depression in ALSP (*Csf1r^+/-^*) Mice. (A-D) Cognitive assessment. The test performed, age of the mice, retention interval (RI) indicated in each panel. The number of mice/genotype in each experiment is shown in bars, in the left panel. (A) Left (training), preference for the left side by *Csf1r^+/-^* mice exploring two familiar identical objects. Right (testing), *Csf1r^+/-^* mice spent less time exploring the novel object (left side). (B) Left, similar exploration of the arms of the Y-maze in all experimental groups. Right, lower discrimination for the novel arm by *Csf1r^+/-^* mice is corrected in *Csf1r^+/-^; Csf2^+/-^ (Dhet*) mice. (C) Left, all experimental groups exhibited similar times of exploration of either object. Right, lower preference for the displaced object by *Csf1r^+/-^* mice is corrected in *Dhet* mice. (D) Left (training), all experimental groups exhibited similar times of exploration of two familiar identical objects. Right (testing), reduced long-term memory for the novel object by *Csf1r^+/-^* mice was corrected in *Dhet* mice. (E) Increased depression-like behavior in male *Csf1r^+/-^* mice is corrected in *Dhet* mice. Data were analyzed using two-way ANOVA followed by Bonferroni’s (A-C), Holm-Sidak’s (D) or Benjamini, Krieger and Yekutieli’s (E) post-hoc tests. The left panel in B was analyzed by one-way ANOVA (not significant). See also Figure S2.

### *Csf2* Heterozygosity Prevents Depression-like Behavior in Male ALSP Mice

Previous studies demonstrated male-specific, depression-like behavior in *Csf1r^+/-^* mice (Chitu et al., 2015). This phenotype was reproduced with males in the current cohort and prevented by *Csf2* heterozygosity (Fig. 2E).

### Monoallelic *Csf2* Inactivation Prevents Olfactory Dysfunction in ALSP Mice

As *Csf1r^+/-^* mice exhibit olfactory deficits (Chitu et al., 2015), we explored the contribution of CSF-2 to this phenotype in an odor discrimination test (Fig. 3A). Mice of all genotypes exhibited lower exploration of the repulsive odorant and trigeminal stimulant, lime, compared with the attractive pure odorant, vanilla. However, in response to vanilla, wild type mice increased their exploration with time, whereas *Csf1r^+/-^* mice failed to do so. This phenotype of *Csf1r^+/-^* mice was corrected by *Csf2* heterozygosity. To further explore the olfactory response of *Csf1r^+/-^* mice to pure odorants, we subjected the mice to an odor threshold assay (Witt et al., 2009), using 2-phenylethanol (Doty et al., 1978) (Fig. 3B). Whereas wild type control mice were able to detect 2-phenylethanol at a dilution of 10^−1^, *Csf1r^+/-^* mice failed to detect the odorant at any concentration. *Csf2* heterozygosity restored detection to *Csf1r^+/-^* mice at a threshold of 10^−2^. These results indicate that the response of *Csf1r^+/-^* mice to pure odorants is impaired and that monoallelic *Csf2* inactivation alleviates this phenotype. *Csf2* heterozygous mice were unable to detect 2-phenylethanol, but responded normally to vanilla. The basis of this selective impairment is unclear.

**Figure 3.**
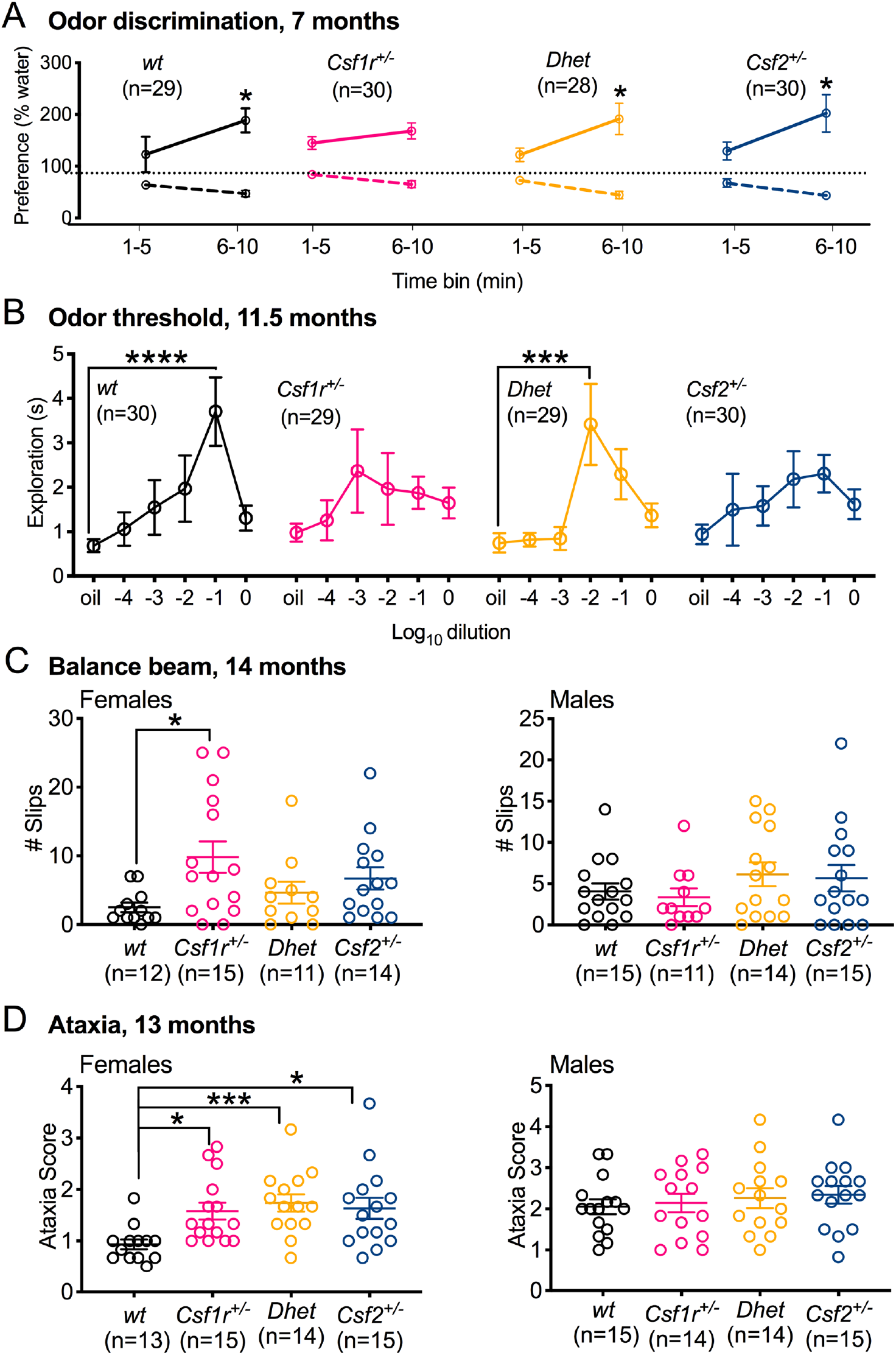
Attenuation of the Olfactory and Motor Coordination Deficits of *Csf1r^+/-^* Mice by *Csf2* Heterozygosity. (A) Odor discrimination at 7 months of age. *Csf1r^+/-^* mice showed no significant increase in exploring the pure odorant vanilla. (B) Odor threshold to the pure odorant 2-phenylethanol by 11.5-month-old mice. Absence of a significant threshold in *Csf1r^+/-^* mice is corrected in *Dhet* mice. (C) Locomotor coordination in mice assessed as number of slips in the balance beam test. (D) Ataxia score in mice assessed as sum of the ledge, hind limb and gait scores. Data were analyzed using two-way ANOVA followed by Bonferroni’s (A) and Dunnett’s (B) post-hoc tests or by Kruskal-Wallis followed by Dunn’s post-hoc tests (C, D).

### Inactivation of a Single *Csf2* Allele Partially Improves the Motor Coordination Deficits of Female *Csf1r^+/-^* Mice

Balance beam studies have shown that older *Csf1r^+/-^* mice possess motor deficits (Chitu et al., 2015). This phenotype was reproduced with female *Csf1r^+/-^* mice. In contrast, double heterozygous mice were indistinguishable from wild type (p = 0.99) (Fig. 3C). Similar to their behavior in the balance beam test, female, but not male *Csf1r^+/-^* mice, exhibited increased ataxia scores (Guyenet et al., 2010) (Fig. 3D) that were not attenuated by *Csf2* heterozygosity. Furthermore, *Csf2* heterozygosity alone produced an ataxic phenotype. Overall, these results show that *Csf1r^+/-^* mice have a female-specific motor deficit and that targeting *Csf2* improves motor coordination on the balance beam, but fails to improve ataxic behavior.

### *Csf2* Heterozygosity Prevents the Cerebral, but not the Cerebellar Microgliosis of *Csf1r^+/-^* Mice

Examination of the effects of inactivation of a single *Csf2* allele in 18-month-old symptomatic *Csf1r^+/-^* mice revealed that *Csf2* heterozygosity prevented the increase in Iba-1^+^ cells in all grey and white matter tracts examined with the exception of the cerebellum (Fig. 4 A, B). A remarkable feature was the presence of periventricular patches of high microglial density in the callosal white matter (Supplemental Fig. S3A, B). Examination of multiple sagittal sections revealed that microglial patches were more frequently encountered in *Csf1r^+/-^* mice than in wild type mice and their frequency was normalized in *Csf1r^+/-^*; *Csf2^+/-^* double heterozygous (*Dhet*) mice (Supplemental Fig. S3B). Morphometric analysis showed that microglia in the white matter patches of *Csf1r+/-* mice had less ramified processes than those of wt mice (Fig. 4 C, D), suggestive of an activated state. The ramified morphology was restored in *Dhet* mice. In contrast, cortical microglia showed no significant difference in ramification compared to wt (Fig. 4 C, E).

**Figure 4.**
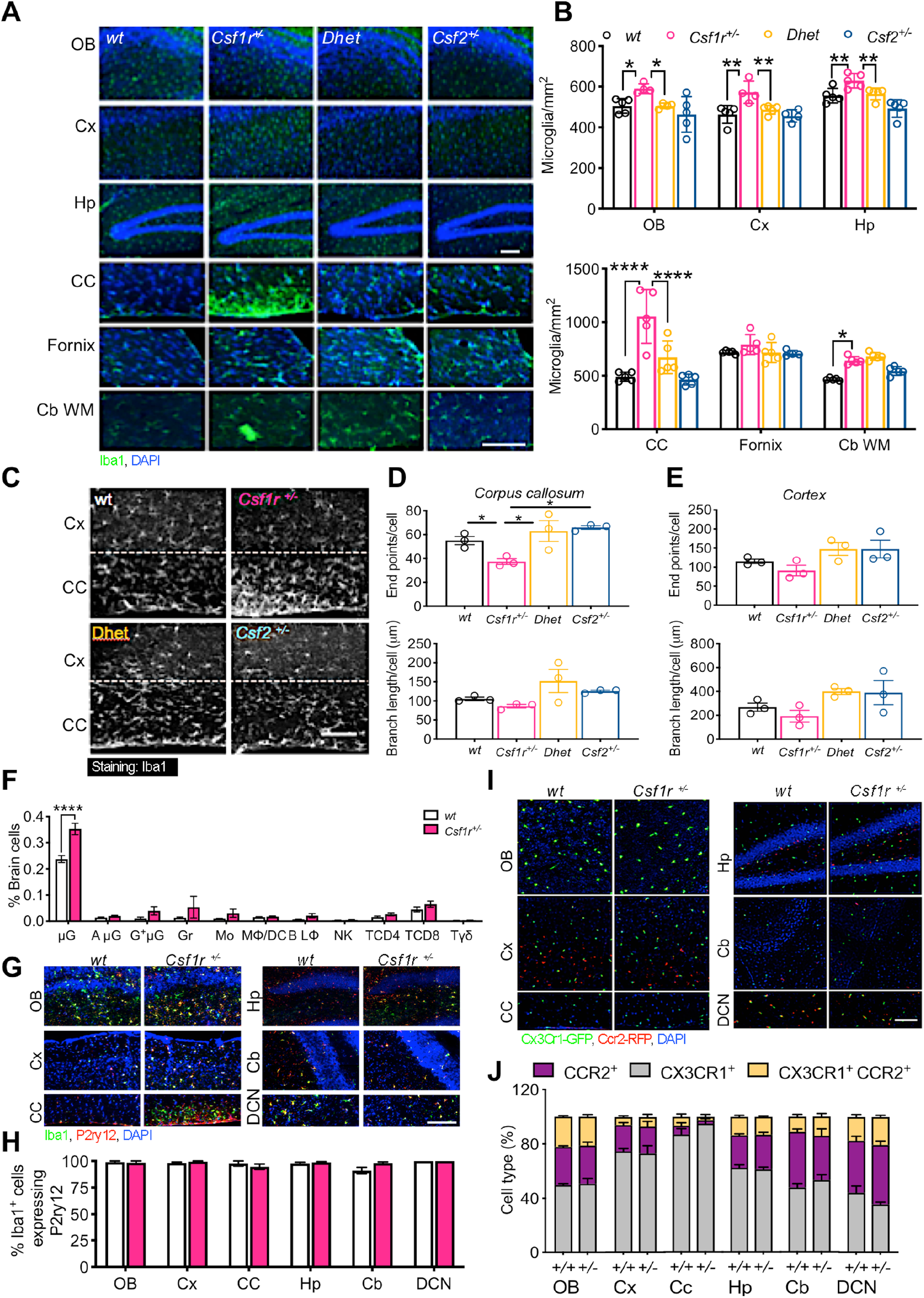
*Csf2* Heterozygosity Prevents Cerebral Microgliosis in aged ALSP Mice. (A) Iba1^+^ cell densities (green) in different areas of the brains of 18-month-old mice. OB, olfactory bulb; Cx, primary motor cortex; CC, corpus callosum; Hp, hippocampus; Cb WM, cerebellar white matter; Cb, cerebellum; DCN, deep cerebellar nuclei. (B) Quantification of Iba1^+^ cell densities. (C) Morphology of Iba1-positive cells. The dotted line indicates the border between the corpus callosum and the adjacent grey matter. (D, E) Quantification of microglia ramification in the white (D) and grey (E) matter regions shown in (C). (F) Quantification of microglia and infiltrating leukocytes by flow cytometry. μG, microglia; AμG, activated microglia; G^+^μG, Ly6G^+^ P2ry12^high^ microglia; Gr, Ly6G^+^P2ry12^-^ granulocytes; Mo, Ly6C^+^ infiltrated monocytes; MΦ/DC, Ly6C^-^ macrophages/dendritic cells; BLΦ, B lymphocytes; NK, natural killer cells; TCD4 and TCD8, CD4 and CD8^+^ T lymphocytes; Tγδ, γδ T cells. Data obtained from 16 month-old, wt (n=4) and *Csf1r^+/-^* (n=5) mice. (G) Colocalization of P2ry12 (red) with Iba1^+^ cells (green). (H) Quantification of P2ry12 expression in Iba1^+^ cells in wild type and *Csf1r^+/-^* mice; 5 mice/genotype. (I) Expression of *Cx3Cr1* (GFP, green) and *Ccr2* (RFP, red) reporters in 11-month-old *Cx3Cr1^GFP/+^;Ccr2^RFP/+^;Csf1r^+/+^* (+/+) and *Cx3Cr1^GFP/+^;Ccr2^RFP/+^;Csf1r^+/-^* (+/-) mice. (J) Quantification of mononuclear phagocytes in *Cx3Cr1^GFP/+^;Ccr2^RFP/+^* reporter mice (5 mice/genotype). Scale bars, 100 μm. Significance was analyzed using two-way ANOVA (B, F, H, J) or one-way ANOVA (D, E) followed by Benjamini Krieger and Yekutieli post-hoc analysis. See also Figures S3 and S4.

### Absence of leukocytic infiltration in the brains of *Csf1r^+/-^* mice

The overexpression of *Csf2* in peripheral helper T cells has been reported to promote monocytic infiltration in the brain (Spath et al., 2017) that could contribute to the expansion of Iba1+ cells (Greter et al., 2015). To address the contribution of peripheral monocytes, we performed an unbiased flow cytometric analysis of all CD45^+^ cells in the brains of 15-month-old mice (Fig 4F, Supplemental Figure S4). This revealed that in both wt and *Csf1r^+/-^* mice, most of the Ly6G^-^ CD45^+^ cells were CD11b^+^ CD45^low^ P2ry12^high^, a profile that identifies brain-resident microglia (Greter et al., 2015, Butovsky et al., 2014). Consistent with this, nearly 100% of Iba1^+^ cells expressed P2ry12 in all brain regions tested (Fig. 4G, H). Analysis of the leukocyte populations revealed that *Csf1r^+/-^* mice did not exhibit increased CD45^high^ CD11b^+^ P2ry12^-^ Ly6C^low^ macrophages/dendritic cells, nor evidence of increased infiltration of P2ry12^-^ Ly6G^+^ granulocytes, CD45^high^ CD11b^+^ P2ry12^-^ Ly6C^hi^ monocytes, or of various lymphocyte populations, compared to wt mice (Fig. 4F). The presence of an unusual population of Ly6G^+^ P2ry12^high^ cells that presumably represent an activated state of microglia (G^+^μG, Fig 4F, Supplemental Figure S4) was also detected in aged mice of both genotypes (p = 0.73 vs. wt). Furthermore, in *Cx3Cr1^GFP/+^; Ccr2^RFP/+^* mononuclear phagocyte reporter mice (Mizutani et al., 2012), regardless of genotype or region, the majority of brain mononuclear phagocytes were GFP single positive and identified as resident microglia (Fig. 4 I, J;). The proportions of RFP (*Ccr2*) single positive monocytes and of cells expressing both monocytic (*Ccr2*) and microglial (P2ry12) markers in *Csf1r^+/-^* mice were comparable to wt (Fig. 4 I, J). These data indicate that there is no increase in leukocytic infiltration in *Csf1r^+/-^* mice. Together with the data shown in Fig. 1D, E, these results indicate that the expansion of Iba1^+^ cells occurs by direct stimulation of resident microglial proliferation by CSF-2.

### Gene Expression Changes in *Csf1r^+/-^* Microglia Suggest a Maladaptive Phenotype

To determine how reductions in *Csf1r* or *Csf2* expression alone, or in combination, affect microglial function in aged (21 month-old) mice, we analyzed the changes in the transcriptome of cerebral Tmem119^+^ microglia compared to wild type controls. *Csf1r* heterozygosity led to the differential expression of 496 genes, comprising 237 upregulated genes (URG) and 259 downregulated genes (DRG) (Supplemental Table 2, Fig. 5A). Functional enrichment analysis revealed that a significant proportion of the *Csf1r^+/-^* URG encoded membrane (57), extracellular (39) and mitochondrial (16) proteins (Supplemental Table 3). In contrast, the innate immunity cluster contained only four URG (*Tgtp1, Ly86* and the complement proteins *C1qb* and *C1qc*). Among the URG encoding extracellular proteins, were transcripts for the secreted inducer of senescence, Augurin (*Ecrg4*) (Kujuro et al., 2010), the neuropeptide Tac2 (Andero et al., 2016) and the CSF-2-induced proinflammatory chemokine Ccl17 (Achuthan et al., 2016) (Fig. 5B). Other upregulated transcripts included those encoding several mitotoxic (*Apoo, Mrps6, Nr2f2, Coq7*) (Turkieh et al., 2014, Sultan et al., 2007, Wu et al., 2015, Lapointe and Hekimi, 2008) and neurodegeneration-related (*Syngrl, Cst7, Trem2, Spp1, Ch25h*) (Hegyi, 2017, Ma et al., 2011, Krasemann et al., 2017, Shin et al., 2011) protein products (Fig. 5B, F, H). Interestingly, *Csf1r* heterozygosity did not reduce the expression of known neurotrophic factors by microglia, rather it enhanced the expression of neurturin, neudesin, midkine, IGF2 and VEGFB (Fig. 5B and Supplemental Table 4), suggesting that microglial neurotrophic functions were not impaired.

**Figure 5.**
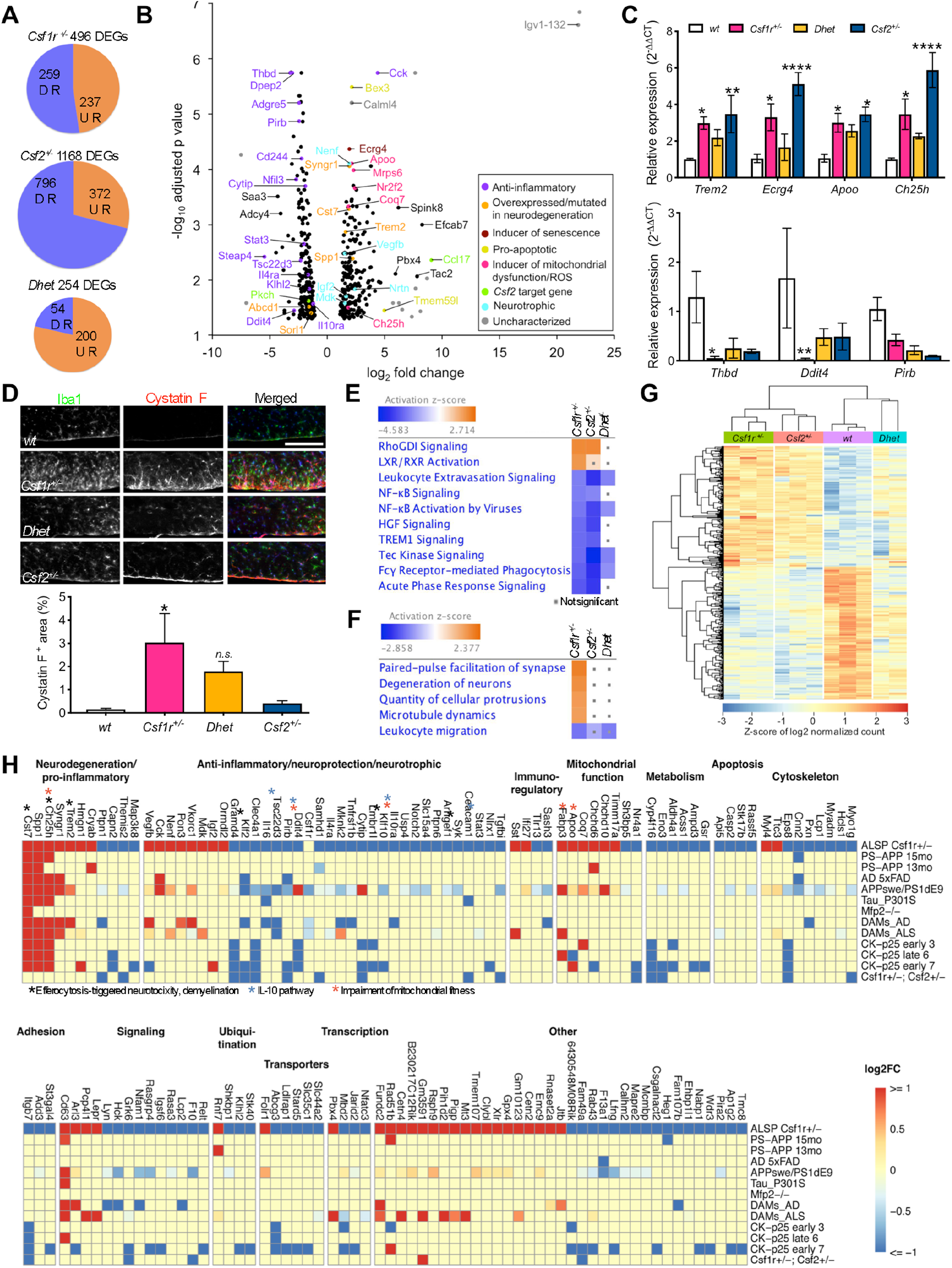
*Csf2* Heterozygosity Restores the *Csf1r^+/-^* Microglial Transcriptomic Profile. (A) Differences in gene expression profile in *Csf1r^+/-^, Csf2^+/-^* and *Dhet* microglia compared to wild type controls. (B) Volcano plot highlighting DEGs of interest in *Csf1r^+/-^* microglia. (C) Validation of changes in expression of selected upregulated (top panel) and downregulated (lower panel) genes inmicroglia isolated from 4 wt, 5 *Csf1r^+/-^*, 5 *Dhet* and 4 *Csf2^+/-^* mice. Two-way ANOVA followed by Dunnett’s post-hoc test. (D) Expression of Cystatin F in the corpus callosum of wt, single and double heterozygous mice. Scale bar, 100 μm. N=5 mice/genotype, One-way ANOVA followed by Tukey’s post-hoc test; n.s., not significant (p =0.33). (E and F) IPA-generated list of pathways (E) and biological processes (F) affected by *Csf1r* heterozygosity and their predicted activation status in *Csf2^+/-^* and *Dhet* microglia. Dots indicate no significant difference. (G) Heatmap showing the expression of *Csf1r^+/-^* DEGs across individual samples. (H) Illustration of the overlap of *Csf1r^+/-^* DEGs with genes differentially expressed in other mouse models of neurodegenerative disease. Note decreased *Csf1r* expression in a model of AD (APPswe/PS1dEp) and in disease-asssociated microglia (DAMs-AD, DAMs-ALS). The *Csf1r* targeting strategy (Dai et al., 2002) does not affect transcription precluding confident detection of decreased *Csf1r* expression in *Csf1r^+/-^* microglia using RNASeq (log2FC = −1.15; p = 0.01; adjusted p value = 0.1).

Analysis of the DRGs showed that approximately 50% (132 out of 259) of these encode membrane proteins (Supplemental Table 3) including proteins with anti-inflammatory activity such as *Thbd, Dpep2, Pirb, Cd244, Il4ra* and *Il10ra* (Wolter et al., 2016, Habib et al., 2003, Zhang et al., 2005, Georgoudaki et al., 2015, Mori et al., 2016, Lobo-Silva et al., 2016) (Fig. 5B, F). Consistent with the downregulation of IL-10 receptor signaling in *Csf1r^+/-^* microglia, its downstream signaling mediator, *Stat3* and several IL-10 transcriptional targets (*Ddit4, Nfil3, Tsc22d3*) (Ip et al., 2017, Lang et al., 2002, Berrebi et al., 2003, Hoppstadter et al., 2015) were also downregulated (Fig 5B, F).

Other potentially relevant downregulated genes encode transcripts associated with Alzheimer’s disease (*Sorl1*) (Nicolas et al., 2016), leukodystrophy (*Abcd1* and its downstream mediator of pathology, *Ch25h*) (Gong et al., 2017, Jang et al., 2016) and the CSF-2 target gene *Pkch*, encoding protein kinase Cη, a regulator of lipid metabolism and suppressor of NO production by macrophages (Torisu et al., 2016, Ozawa et al., 2016) (Fig 5B).

Pathway analysis revealed that the transcriptomic changes associated with *Csf1r* heterozygosity are consistent with activation of the RhoGDI signaling and LXR/RXR pathways (Fig. 5E, Supplemental Table 5). The LXR/RXR pathway has been reported to increase cholesterol efflux and repress TLR4-induced genes (Hiebl et al., 2018), but also to promote inflammasome activation in microglia (Jang et al., 2016). However, inhibition of classical pro-inflammatory pathways, such as the acute phase response and NFκB signaling, as well as of TREM1 signaling, that sustains inflammation (Owens et al., 2017), was also predicted (Fig. 5E and Supplemental Table 5), suggesting that *Csf1r^+/-^* microglia are not pro-inflammatory. Analysis of the biological processes affected by *Csf1r* heterozygosity predicted active neurodegeneration, increased cellular protrusions and microtubule dynamics, as well as an elevated paired-pulse facilitation of synapses (Fig. 5F and Supplemental Table 6).

To further explore how these transcriptomic changes are relevant to the neuropathology, we intersected our gene list with publicly available datasets showing changes in the microglial transcriptome in other mouse models of neurodegenerative disease, including models of Alzheimer’s disease (PS-APP, AD 5xFAD, APPswe/PS1dE9), tauopathy (Tau_P301S), amyotrophic lateral sclerosis (ALS), rapid neurodegeneration (CK-p25) and spinocerebellar ataxia (*Mfp2^-/-^*) (Friedman et al., 2018, Keren-Shaul et al., 2017, Mathys et al., 2017) (Fig 5H). A list showing the genes similarly regulated and their functions is provided in Supplemental Table 7. The comparison shows that 29% of the *Csf1r^+/-^* DEGs were similarly regulated in at least one other neurodegenerative disease, the most extensive overlap occurring with the APPswe/PS1dE9 Alzheimer’s disease model, followed by changes related to early neurodegeneration (CK-p25 early cluster 7) and to ALS. Consistent with this overlap, *Csf1r* expression was downregulated in microglia isolated from mouse models of Alzheimer’s disease and ALS (Fig. 5H). Changes occurring in most neurodegenerative conditions were the high expression of a group of transcripts encoding the lysososmal cathepsin inhibitor Cystatin F (*Cst7*) (Ma et al., 2011), osteopontin (*Spp1*), an opsonin for cell debris (Shin et al., 2011), and cholesterol 25-hydroxylase (*Ch25h*) that, through its product, 25-hydroxycholesterol, activates the LXR pathway and promotes ROS production and inflammation (Jang et al., 2016). Underexpression of neuroprotective (*Clec4a1, Il16*) (Flytzani et al., 2013, Shrestha et al., 2014) and anti-inflammatory (*Klf2, Gramd4, Ddit4, Pirb, Tsc22d3*) (Roberts et al., 2017, Ip et al., 2017, Kimura et al., 2015, Zhang et al., 2005, Berrebi et al., 2003) transcripts was observed in ALSP and at least three other conditions (Fig. 5H).

Together, our data suggest that *Csf1r* heterozygosity does not produce a neurotrophic defect, or an overt inflammatory activation of microglia. Rather, the transcriptional profile predicts a maladaptive microglial phenotype (Fig. 6A). A summary of the pathways that could contribute to disease pathology is presented in Fig. 6B.

**Figure 6.**
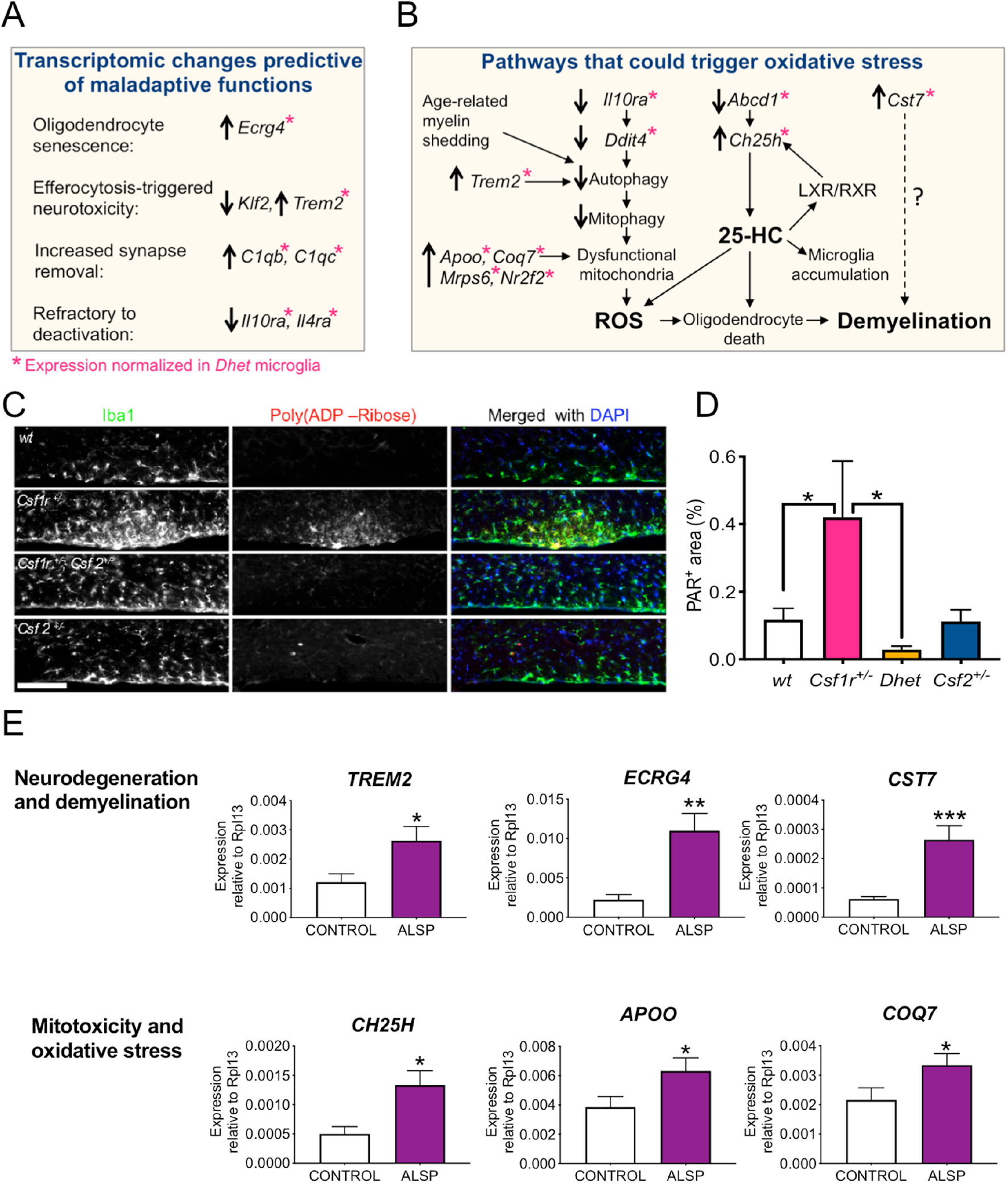
Pathways dysregulated in *Csf1r^+/-^* mice and ALSP patients. (A, B) Predicted madalaptive functions (A) and hypothetical pathways (B) dysregulated in *Csf1r^+/-^* mouse microglia (C) Evidence of oxidative stress: colocalization of the poly (ADP-Ribose) signal with callosal microglial patches in the periventricular white matter. Scale bar, 100 μm. (D) Quantification of the callosal area positive for poly (ADP-Ribose) in 2-3 sections/mouse, 5-9 mice/genotype. One-way ANOVA followed by Kruskal-Wallis test. (E) Transcriptomic changes potentially critical for pathology also occur in the periventricular white matter of ALSP patients and healthy controls (n=5); *, p < 0.05, one-tailed Student’s t test.

### *Csf2* Heterozygosity Upregulates Antioxidant and Anti-inflammatory Signaling in Microglia

As described above, *Csf2* insufficiency alone also impaired cognition (Fig. 2) and partially affected olfaction and motor coordination (Fig. 3). However, these phenotypes were not accompanied by microgliosis in *Csf2^+/-^* mice. Thus, we examined the effect of *Csf2* heterozygosity on microglia. Compared to wt controls, *Csf2* heterozygosity dysregulated the expression of 1168 genes (372 URG and 796 DRG) (Fig. 5A and Supplemental Table 1). Most of the top differentially expressed transcripts encode products yet uncharacterized or pseudogene transcripts (Supplemental Fig. S5A). Among the top upregulated transcripts were those encoding the proinflammatory cytokine CCL3 and the matrix metalloproteinase MMP11. The myelin-degrading MMP12 (Hansmann et al., 2012) was also upregulated (Supplemental Fig. S5D). Like *Csf1r^+/-^* microglia, *Csf2^+/-^* microglia downregulated the expression of several anti-inflammatory genes (Supplemental Fig. S5A and D). Remarkably, *Csf2^+/-^* microglia reduced the expression of *Il4ra*, but not of *Il10ra* (Supplemental Fig. S5D and Supplemental Tables 2 and 8).

Analysis of pathways uniquely affected by *Csf2* heterozygosity showed activation of the antioxidant vitamin C pathway, of PPAR signaling, which regulates lipid metabolism and is anti-inflammatory (Wahli and Michalik, 2012) and of the complement system that has an well-established role in synapse loss in neurodegenerative disease (Hajishengallis et al., 2017, Stephan et al., 2012). Among the top predictions for inhibited pathways, we found dendritic cell maturation, neuroinflammation, LPS and iNOS signaling (Supplemental Fig. S5B). Analysis of biological processes selectively affected by *Csf2* heterozygosity predicted increased inflammation and encephalitis that were paradoxically associated with decreased leukocyte infiltration, as well as decreased activation of neuroglia and microglia (Supplemental Fig. S5C). These data suggest that the neuropathology in *Csf2^+/-^* mice is not associated with classically defined inflammatory activation (i.e. increased proinflammatory cytokine and iNOS expression). CSF-2 insufficiency increases the activation of the complement system and the expression of MMPs, both of which could affect neuronal network structure and function. However, it also causes the activation of antioxidant (Vitamin C antioxidant pathway, upregulation of *Prdx4*) and lipid metabolic (PPAR signaling) pathways that are expected to reduce oxidative stress and inflammation.

### Targeting *Csf2* in ALSP Mice Attenuates Microglial Dysfunction and Oxidative Stress

Investigation of the effect of reduction of CSF-2 availability revealed a large decrease in the microglial transcriptomic changes of *Csf1r^+/-^;Csf2^+/-^* microglia (254 DEGs, 54 URG and 200 DRG) compared to *Csf1r^+/-^* microglia (Fig. 5A). Indeed, hierarchical clustering of samples based on DEGs grouped *Csf1r^+/-^* and *Csf2^+/-^* apart from the double heterozygous samples, which were more related to wild type samples (Fig. 5G). Consistent with this, pathway analysis predicted the restoration of RhoGDI and LXR/RXR signaling and attenuation of the neurodegenerative phenotype (Fig. 5 E, F). Furthermore, 86% of the transcriptional changes common to ALSP and other neurodegenerative conditions were eliminated in *Csf1r^+/-^*;*Csf2^+/-^* microglia (Fig. 5H), including the expression of gene products that suppress mitochondrial fitness and enhance oxidative stress (e.g. *Ch25h, Ddit4, Il10ra, Apoo, and Coq7*). Consistent with this, monoallelic inactivation of *Csf2* in *Csf1^+/-^* mice reduced poly ADP-ribosylation, a marker of oxidative stress, in the periventricular white matter microglial patches (Fig. 6C, D).

The *Cst7*-encoded protein, Cystatin F, is a microglial marker of ongoing demyelination with concurrent remyelination (Ma et al., 2011). As predicted by the changes in *Cst7* transcript abundance, Cystatin F protein was readily detected in the callosal microglial patches present in *Csf1r^+/-^* brains, but its staining in double heterozygous brains was not significantly different from the wild type staining (Fig. 5D). Thus, targeting CSF-2 improves microglial homeostatic functions.

### Increased Expression of CSF-2 target Genes Potentially Relevant to Pathology in the White Matter of ALSP Patients

To determine whether similar gene expression changes occur in ALSP patients, we isolated RNA from the callosal white matter of ALSP patients and control (Supplemental Table 1) brains and performed real time qPCR. Several transcripts potentially relevant to neurodegeneration, demyelination and oxidative stress were also upregulated in the white matter of ALSP patients (Fig. 6E). These results identify putative common contributors to the pathology of ALSP and of other demyelinating and neurodegenerative diseases.

### Monoallelic *Csf2* Inactivation on the ALSP Background Improves Callosal Myelination

Reduction of the expression of the demyelination marker, Cystatin F, by targeting *Csf2*, prompted us to examine the ultrastructure of the corpora callosa of all genotypes by transmission electron microscopy. Examination of cross-sections showed that *Csf1r^+/-^* fibers have higher G-ratios than wild type fibers, indicative of demyelination followed by remyelination (Fig. 7 A-C). Higher G-ratios were also observed in *Csf2^+/-^* samples (Fig. 7A, B, E). The G-ratios of callosal fibers in double heterozygous mice were not significantly different from those of wild type fibers (Fig. 7A, B, D). Consistent with this, the staining for myelin basic protein (MBP) was reduced in the corpus callosum of *Csf1r^+/-^* mice and the reduction prevented by targeting *Csf2* (Fig. 7F). Other white matter tracts (fimbria and the cerebellar white matter) although trending similarly, were not significantly affected (Fig. 7F). Changes in myelination did not result from decreased availability of PDGFRα^+^ early oligodendrocyte precursors or of mature CC1^+^ oligodendrocytes, which were paradoxically increased (Fig. 7G). Examination of age-related changes in myelin compaction revealed significantly increased myelin degeneration in the *Csf1r^+/-^* mice that was not rescued by *Csf2* heterozygosity (Fig. 7H). In terms of axonal pathology the data reflect a lack of protective effect of *Csf2* targeting on neurodegeneration (Fig. 7I). Consistent with the overall lack of protection against neurodegeneration, *Csf2* heterozygosity did not attenuate the loss of NeuN^+^ mature neurons in cortical layer V (Fig 7J, K). Together, these data indicate that *Csf2* heterozygosity rescues myelination, but is not sufficient to prevent neurodegeneration and the exacerbation of age-related myelin degeneration in ALSP mice.

**Figure 7.**
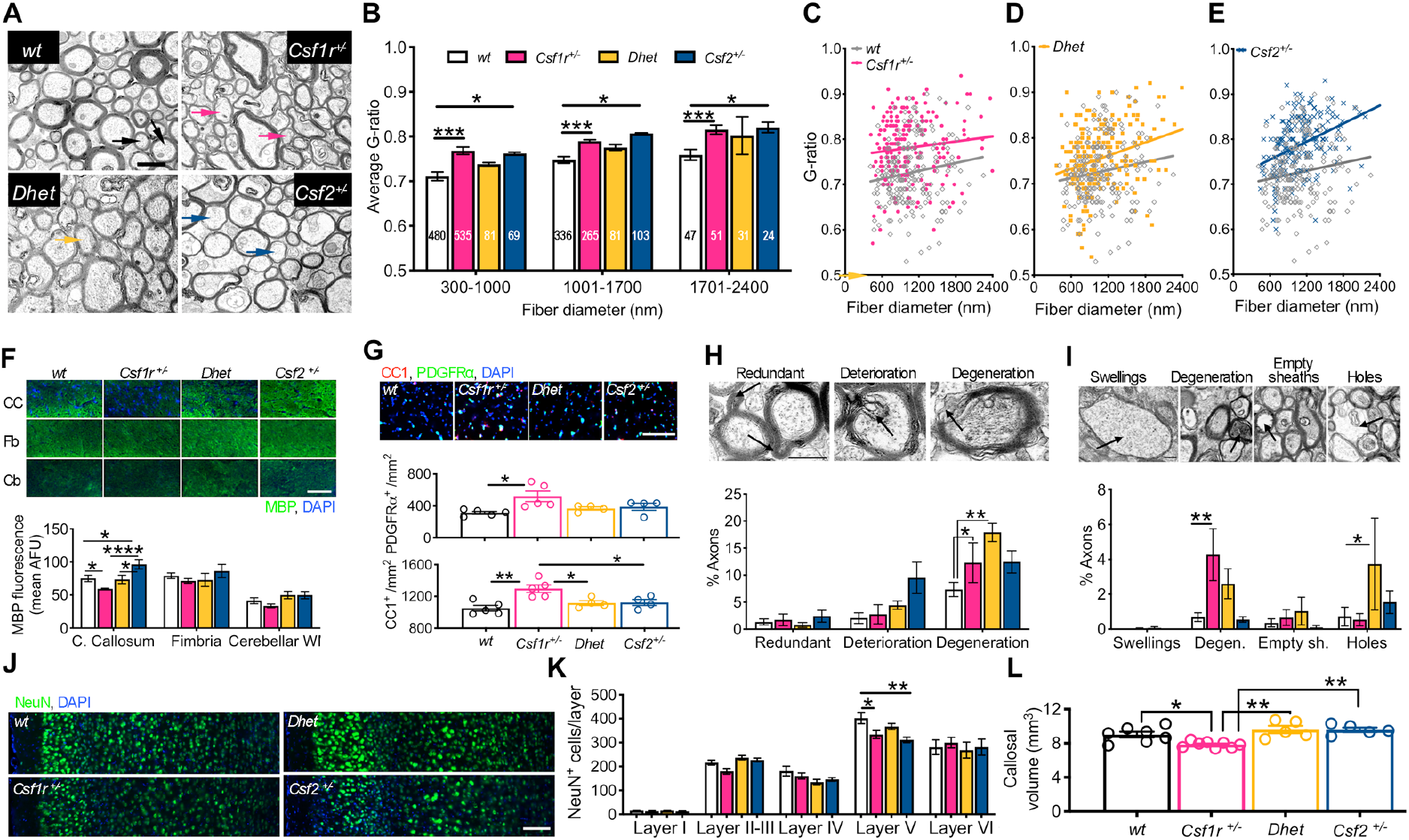
*Csf2* Heterozygosity in ALSP Mice Prevents Callosal Atrophy and Improves Myelination. (A) Myelin and axonal ultrastructure in callosal cross-sections from 9-11-month-old mice. Arrows point to examples of changes in myelin thickness in axons of small and medium diameters. (B-E) Changes in G-ratio in *Csf1r^+/-^* (B, C) and *Csf2+/-* (B, E) mice are attenuated by double heterozygosity (B, D). Panel B shows average values per mouse (2-6 mice/genotype); values in the bars indicate the total numbers of fibers examined in each fiber diameter range. Panels C-E show individual G-ratio values. (F) Quantification of MBP staining in white matter tracts including corpus callosum (CC), fimbria (Fb) and cerebellum (Cb). N= 3-7 mice/genotype. (G) Changes in myelination are not accompanied by a decrease in early oligodendrocyte precursors (PDGFRα^+^) or oligodendrocytes (CC1^+^). (H) Quantification of age-induced myelin pathology in wt and mutant mice (2-7 mice/genotype, > 900 neurons/genotype). The upper panels show representative examples of structural abnormalities. (I) Quantification of age-induced axonal pathology in wt and mutant mice (data from 3-6 mice/genotype, > 900 neurons/genotype). The upper panels show representative examples of structural abnormalities. (J) Neuronal loss in cortical layer V at 18 months of age. Scale bar, 100μm. (K) Average NeuN positive cells/layer. N= 4 mice/genotype. (L) *Csf2* heterozygosity prevents callosal atrophy in 19-month old *Csf1r^+/-^* mice. Significance was analyzed using two-way ANOVA followed by Holm-Sidak’s (B, F, K) or Benjamini, Kreiger and Yekutieli’s (H, I) post-hoc tests and one-way ANOVA followed by Tukey’s post-hoc test (H, L). Scale bars: 1μm in (A), 50μm in (F), 100μm in (G) and (J), 500nm in (H) and (I).

### *Csf2* Heterozygosity Normalizes the Callosal Volume in *Csf1r^+/-^* Mice

Since *Csf2* heterozygosity in the ALSP background normalized the G-ratios, we examined whether this resulted in attenuation of white matter loss, using MRI. Compared with wild type, callosal volumes were lower in *Csf1r^+/-^* mice and *Csf2* heterozygosity prevented callosal atrophy (Fig. 7L). In contrast, *Csf2* heterozygosity did not cause a reduction in callosal volume.

## Discussion

In a previous study (Chitu et al., 2015) we showed that microglial densities were elevated in several brain regions of young and old *Csf1r^+/-^* ALSP mice and associated with increased expression of *Csf2*, encoding the microglial mitogen, CSF-2. Aside from its mitogenic activity, CSF-2 primes neurotoxic (Fischer et al., 1993) and demyelinating (Smith, 1993) responses in microglia. As *CSF2* expression is also elevated in post-mortem ALSP brains (Fig. 1F), we reasoned that CSF-2 plays an important role in ALSP pathogenesis. Indeed, we show that *Csf2* heterozygosity rescues the olfactory, cognitive and depression-like phenotypes of *Csf1r^+/-^* mice and ameliorates the motor coordination deficits (Figs. 2–3). *Csf2* targeting also reduces the microgliosis (Figs 1A,B and 4A,B) and a hallmark feature of ALSP, demyelination (Fig. 7A-D, F). Although in the CNS, the CSF-2 receptor is expressed both on neural lineage cells (Reed et al., 2005, Baldwin et al., 1993) and microglia, several lines of evidence suggest that *Csf2* deletion ameliorates neuropathology by acting on microglia, rather than on neural lineage cells. First, the reported neuroprotective (Schabitz et al., 2008) and oligodendrogenic (Baldwin et al., 1993) activities of CSF-2 suggest that its targeting in ALSP should be detrimental, rather than protective. However, we found that *Csf2* heterozygosity did not exacerbate the loss of Layer V neurons, indicating that its neuroprotective actions are negligible in the context of ALSP. Furthermore, in the ALSP mouse, demyelination is paradoxically associated with an expansion of oligodendrocyte precursors and APC^+^ oligodendrocytes (Fig. 7G). Targeting of *Csf2* prevents the increase in oligodendrocytes and oligodendrocyte precursors in ALSP mice (Fig. 7G), yet this finding cannot explain how it attenuates the loss of myelin. One conceivable explanation is that *Csf2* heterozygosity prevents microgliosis (Lee et al., 1994) and the priming of demyelinating (Smith, 1993) and neurotoxic (Fischer et al., 1993) responses in microglia. A direct investigation of the contribution of CSF-2 signaling in different cell types to pathology requires conditional targeting of its receptor, *Csf2ra*, which is presently not possible due to the absence of a specific genetic model. An acceptable approximation is the conditional targeting of *Csf2rb*, which encodes the common subunit of CSF-2, IL-3 and IL-5 receptors (Croxford et al., 2015), using lineage-specific *Cre* drivers. Using this approach, we showed that attenuation of CSF-2 signaling in CX3CR1-expressing cells (i.e. mononuclear phagocytes and microglia) was sufficient to prevent white matter microgliosis in young mice (Fig. 1 D, E). Based on this observation and on the finding that monocyte-derived cells do not significantly contribute to the widespread microgliosis in aged *Csf1r^+/-^* mice (Fig. 4F-J) we conclude that CSF-2 triggers microgliosis *via* direct signaling in CNS-resident microglia. Thus, while a contribution of CSF-2 signaling in other cell types cannot be formally excluded, current data suggests that the CSF-2-mediated dysregulation of microglial function plays a central role in the pathology of ALSP mice. Consistent with this, transcriptomic analysis revealed that *Csf2* heterozygosity suppressed a high proportion of the transcriptomic changes occurring in *Csf1r^+/-^* microglia, including the expression markers of oxidative stress and demyelination (Figs. 5 and 6 and more detailed discussion, below).

Transcriptomic analysis suggests that clearance of apoptotic cells and myelin debris triggers maladaptive responses in *Csf1r^+/-^* microglia. Relevant changes include the overexpression of TREM2, an innate immune receptor expressed on microglia that promotes the clearance of apoptotic neurons and myelin debris (Poliani et al., 2015). Recent work indicates that following the uptake of the apoptotic neurons or neuronal debris, TREM2 increases the expression of oxidative stress markers and complement components and suppresses the homeostatic function of microglia (Linnartz-Gerlach et al., 2019, Krasemann et al., 2017). The molecular mechanism could involve suppression of mitophagy either directly (Ulland et al., 2017, Wang et al., 2019) or as an indirect consequence of overloading of the degradative pathway by the ingested myelin (Safaiyan et al., 2016). Furthermore, as observed in the mouse model (Fig. 5C), overexpression of *TREM2* also occurs in the white matter of ALSP patients (Fig. 6E), where others have documented the presence of lipid-laden macrophages (Tada et al., 2016, Lin et al., 2010). Thus, decreased autophagy may contribute to ALSP pathology and the benefits of stimulation of autophagy should be further explored.

Dysregulation of lipid metabolism may also contribute to ALSP. One of the genes downregulated in *Csf1r^+/-^* microglia encodes the very long chain fatty acid transporter ABCD1. Mutations in *ABCD1* cause X-linked adrenoleukodystrophy, a demyelinating disease associated with microglial dysfunction mediated by the overexpression of *Ch25h* encoding cholesterol 25 hydroxylase (Gong et al., 2017, Jang et al., 2016). The product of CH25H, 25-hydroxycholesterol (25-HC), is an activating ligand of LXR providing an explanation for the activation of the LXR/RXR pathway in *Csf1r^+/-^* microglia (Fig. 5E). In vivo, 25-HC was reported to promote oligodendrocyte death and, *in vitro*, 25-HC stimulates IL-1β secretion by microglia in a mitochondrial ROS- and LXR/RXR-dependent manner (Jang et al., 2016). The expression of *CH25H* was also elevated in the white matter of ALSP patients (Fig. 6E), suggesting that dysregulation of cholesterol metabolism in microglia may contribute to the pathology of ALSP.

Double *Csf1r* and *Csf2* heterozygosity eliminates the changes in most canonical pathways and biological processes produced by either *Csf1r* or *Csf2* heterozygosity alone, (Fig. 5E, F and Supplemental Fig. S5B, C) including the activation of the LXR/RXR pathway (Fig. 5E). In the context of ALSP, *Csf2* heterozygosity virtually restores microglial function, resulting in attenuation of oxidative stress, improvement of callosal myelin thickness and restoration of the callosal volume (Figs. 5–7). Together with the normalization of most behavioral phenotypes of *Csf1r^+/-^* mice by monoallelic targeting of *Csf2*, these studies clearly identify CSF-2 as a therapeutic target in ALSP. In addition, this work demonstrates that reduction of either CSF-1R, or CSF-2 signaling, impairs microglia function and the homeostasis of the aging CNS and that rebalancing the signals is beneficial. Both increased *CSF2* levels and decreased microglial *Csf1r* expression have been reported in Alzheimer’s disease (Fig. 5) (Tarkowski et al., 2001) and multiple sclerosis (Kostic et al., 2018, Werner et al., 2002). Thus, apart from ALSP, the unbalanced CSF-1R/CSF-2 signaling described here may contribute to the pathogenesis of other neurodegenerative conditions.

## Acknowledgements

We thank Dr. Ian Willis of the Department of Biochemistry for critically evaluating the manuscript, Dr. Shahina B. Maqbool of the Epigenomics Shared Facility for the RNA-Seq, Leslie Cummins and Frank Macaluso of the Analytical Imaging Facility for the preparation of transmission EM samples, Drs. Craig A. Branch and Min-Hui Cui, of the Gruss Magnetic Resonance Research Center for MRI imaging, Hillary Guzik, Andrea Briceno and Dr. Vera DesMarais of the Analytical Imaging Facility for help with imaging and histomorphometry and Drs. Fabien Delahaye and Xusheng Zhang of the Computational Genomics Core Facility for data analysis, all at the Albert Einstein College of Medicine. This work was supported by grants from the National Institutes of Health Grant R01NS091519 (to ERS), U54 HD090260 (support for the Rose F. Kennedy IDDRC), the P30CA013330 NCI Cancer Center Grant and grants from the Swiss National Science Foundation 310030_146130 and 316030_150768 (to B.B.) and PP00P3_144781.

## Author Contributions

VC & ERS designed the study and wrote the manuscript. FB, GGLS, ESP, MEG, VC, KS performed the behavioral experiments. VC, FB, SG, GGLS, HCK carried out the histological and ultrastructural studies. VC, PW, DZ performed the microglial transcriptomic analyses. DS performed the flow cytometric analysis. DWD, MAD collected and prepared the post-mortem brain tissue, DS executed the flow cytometry and VC performed human gene expression analysis. BB and ALC provided the *Csf2rb^fl/fl^* mice. MEG, SG, MFM, ZKW advised in specific areas.

## Methods

### LEAD CONTACT AND MATERIALS AVAILABILITY

Further information and requests for resources and reagents should be directed to and will be fulfilled by Dr. E. Richard Stanley. The study did not generate new unique reagents.

### EXPERMENTAL MODEL AND SUBJECT DETAILS

#### In vivo animal studies

##### Mouse Strains, Breeding and Maintenance

All *in vivo* experiments were conducted in accordance with the National Institutes of Health regulations on the care and use of experimental animals and approved by the Albert Einstein College of Medicine Institutional Animal Care and Use Committee. The generation, maintenance and genotyping of *Csf1^+/-^* mice was described previously (Dai et al., 2002). *Csf2^+/-^* mice (Dranoff et al., 1994) were a gift from Dr. Glenn Dranoff and were genotyped using a PCR procedure developed by the Dranoff laboratory that utilizes the primers listed in the Key Resources table. Both lines were backcrossed for more than 10 generations onto the C57BL6/J background. Cohorts were developed from the progeny of matings of *Csf1r^+/-^* to *Csf2^+/-^* mice, randomized with respect to the litter of origin. At 3 months of age, they were transferred from a breeder diet (PicoLab Rodent Diet 20 5058) to a lower fat maintenance diet (PicoLab Rodent Diet 20 5053). This prevented the increase in body weight in *Csf1r^+/-^* mice compared with wild type mice observed in our earlier study (Chitu et al., 2015) and was also associated with delayed onset of spatial memory deficits (from 7 to 14 months of age) and absence of motor impairment in male *Csf1r^+/-^* mice. *Csf2rb^fl/fl^* mice (Croxford et al., 2015) were provided by Dr. Burkhard Becher, via Dr. William R Drobyski, Medical College of Wisconsin, *Cx3Cr1^GFP/+^; Ccr2^RFP/+^* mononuclear phagocyte reporter mice (Saederup et al., 2010) were a gift from Dr. Susanna Rosi, Kavli Institute for Fundamental Neuroscience, University of California, San Francisco and *Cx3Cr1^Cre/+^* mice (Yona et al., 2013) were a gift from Dr. Marco Prinz, Institute of Neuropathology, Freiburg University Medical Centre, Freiburg, Germany. The age and sex of animals used in each experiment is indicated in the figures.

#### Human studies

Frozen brain tissue blocks containing periventricular white matter were obtained from the Mayo Clinic Brain Bank. Consent for autopsy was obtained from the legal next-of-kin. Studies involving autopsy tissue are exempt from human subjects research (Health and Human Services Regulation 45 CFR Part 46). Information on the ALSP patients harboring *CSF1R* mutations and control cases included in this study is summarized in Supplemental Table 1.

### METHOD DETAILS

#### Cognitive Assessment

Behavioral studies were carried out by blinded operators. Recognition memory was assessed as time exploring a familiar and a novel object, using the novel object recognition test (Ennaceur and Delacour, 1988). The experimental groups were first tested at 7 months of age for short-term memory (1h-retention interval), then at the age of 16 months for long-term memory (24h-retention interval). Mice explored two identical objects (familiarization) for 4 min (1h-retention) or for 10 min (24h-retention) during the training stage. Mice were then exposed to one of the familiar objects and to a novel object for 3 min (1h-retention) or 5 min (24h-retention) during the testing stage. Each mouse was placed in the center of a 40 cm x 40 cm open field box and allowed to explore the objects freely during each stage. Two different pairs of non-toxic objects were used for each experiment. The novelty of the objects (i.e., novel vs. familiar) was counter-balanced within each genotype and the objects were previously validated for equivalent exploratory valence.

Spatial recognition memory was measured at 13.5 months of age in the two-trial test version of the Y-maze (Biundo et al., 2016). Briefly, during the training trial, one of the arms of the maze was closed, and mice were placed into one of the two remaining arms of the maze (start arm) and allowed to explore the open two arms for 10 min. After a 1h inter-trial interval, the blocked arm was opened (novel arm), and mice were placed in the start arm and allowed to explore freely all three arms of the maze for 5 min (test trial).

Spatial recognition memory was also tested at 14.5 months of age in the object placement test (Ennaceur and Delacour, 1988). Each mouse was exposed for 7 min to two identical objects placed in a 40 cm x 40 cm open field box. Four different visual cues were hung on the walls of the box to permit each mouse to orient within the arena. During the training stage, the objects were placed at a distance of 10 cm from one another. After an interval of 25 minutes, one of the objects was displaced into a novel position (15 cm distance, 90 degree angle) and each mouse was returned to the same box to explore the objects for 5 min (testing).

#### Olfactory Assessment

Olfactory discrimination was tested at 8 months of age. Each mouse was exposed to two non-social odors, a pure odorant attractant (vanilla extract) and an aversive odorant (lime extract), and to water as control. Each odorant was adsorbed in a filter paper placed in a 30-mm Petri dish (5 ul of lime extract, 30 ul of vanilla extract, or 30 ul of water per filter). During the test, a single mouse was placed in the center of a 40 cm x 40 cm box in which the dishes containing odorant filters were placed in two opposite corners of the box, while two dishes containing water-adsorbed-filters were placed in the remaining opposite corners. All the dishes were covered until each mouse was placed in the box. Mice were allowed to explore for 10 minutes. Time exploring each odorant and water was recorded during two 5-minute consecutive bins.

The ability of mice to explore a pure odorant was further assessed in 10-month-old mice using the olfactory threshold test in which exploration of the pure odorant, 2-phenylethanol, or mineral oil carrier, applied to a cotton tip, is assessed (Doty et al., 1978, Witt et al., 2009). All experiments were carried in a plexiglas box placed in a odor-free ventilated hood. Before testing, each mouse was habituated to the cotton tip by being exposed five times to mineral oil. During the testing trials, the experimental groups were exposed to 5 ten-fold serial dilutions (from 10^−4^ to 10^0^) of the odorant. Between each dilution of odorant mice were exposed to mineral oil. In each condition, the cumulative time spent exploring the odorant or mineral oil over a 1 minute period was recorded.

#### Depression-like Behavior

Depression-like behavior was assessed as immobility, using the Porsolt Forced Swim Test (Porsolt et al., 1977a, Porsolt et al., 1977b). Briefly, each mouse was placed into a 4-liter beaker filled with warm water (24°C) for 10 minutes and the duration of immobility during three, 3-minute, consecutive bins was recorded.

#### Motor Coordination and Ataxic Behavior

Motor coordination was assessed as the number of slips made while crossing a round, wooden balance beam (Gulinello et al., 2008). Briefly, each mouse was allowed to walk along a 1.6 cm diameter, 1 m long beam placed between two holders 1 meter off the ground. Palatable food was placed at the end of the beam as an incentive to cross. The ataxia phenotype was evaluated as the sum of scores in the ledge, the hindlimb clasping and the gait tests, as previously described (Guyenet et al., 2010).

#### Gene Expression Studies in ALSP Patients

RNA was isolated from either the periventricular white matter or the adjacent gray matter of 5 ALSP patients and 5 control patients (see Supplemntal Table 1) using Trizol and cDNA was prepared using a Super Script III First Strand Synthesis kit (Invitrogen, Carlsbad, CA). Real time PCR was performed using SYBR Green in an Eppendorf Realplex II thermocycler. The primers used are listed in the Key Resources table. Average values from two different blocks of tissue per patient, were used to construct the figures.

#### Analysis of Microglia and Leukocytes

Microglia and brain leukocytes were analyzed using and adaptation of the protocol described by Legroux et al. (Legroux et al., 2015). Briefly, mice were perfused with ice-cold PBS containing 10 U/ml heparin. Brains were dissected, minced and digested with Collagenase D, for 20’at 37°C. Myelin was removed by centrifugation in 37% Percoll. The cells were stained using the antibodies listed in Key Resources Table and analyzed by FACS in a MoFlo Astrios EQ (Beckman Coulter, IN) with a 70-m nozzle. The antibodies used for staining are listed in the Key Resources table and the gating strategy utilized to identify each cell type is shown in Supplemental Figure S3.

#### Microglia Isolation and RNA-Seq Analysis

Microglia were isolated by FACS (Bennett et al., 2016). The RNA was extracted using a Qiagen RNeasy Plus Micro kit and stored at −80°C prior to analysis. We obtained 150 bp paired-end RNA-Seq reads from an Illumina NextSeq 500 instrument. For 3 biological replicates of wild type, *Csf1r*^+/-^, *Csf2^+/-^*, and 2 biological replicates for *Csf1r^+/-^;Csf2^+/-^* samples, an average of ~46 million pairs of reads per sample was obtained. The computational pipeline for identifying differentially expressed genes (DEGs) has been described previously (Wang et al., 2017). Briefly, the Kallisto (v0.43.1) software (Bray et al., 2016) was employed to determine the read count and transcripts per kilobase million (TPM) for each gene that was annotated in the GENCODE database (vM15) (Mudge and Harrow, 2015). To identify DEGs, 14,739 expressed genes with an average TPM >1 were selected in any of the wild type, *Csf1r*^+/-^, *Csf2^+/-^* and *Csf1r^+/-^*;*Csf2^+/-^* samples, using the software DESeq2 (Mudge and Harrow, 2015) and false discovery rate (FDR) < 0.05. For selected genes changes in expression were validated by qPCR, utilizing the primers listed in the Key Resources table.

#### Comparison of DEGs With Other Datasets

The DEG lists from *Csf1r^+/-^* or *Csf2^+/-^* samples were compared to the DEG lists generated from other studies of microglia transcriptome changes associated with neurodegeneration (Mathys et al., 2017, Keren-Shaul et al., 2017). In Keren-Shaul’s (Keren-Shaul et al., 2017) and Friedman’s (Friedman et al., 2018) studies, DEGs were defined as FDR < 0.05, as described in the original papers, while in Mathys’ study (Mathys et al., 2017) DEGs were defined by |z| > 2. The log2(fold-change) of those DEGs were used to generate heatmaps.

#### MRI Imaging

Mice were imaged on an Agilent Direct Drive 9.4 T MRI system (Agilent Technologies, Santa Clara, CA) as previously described (Chitu et al., 2015). Callosal volumes were measured using MIPAV 7.1.1 freeware (mipav.cit.nih.gov).

#### Ultrastructural Studies

Callosal sections were obtained as described (Chitu et al., 2015) and examined by transmission electron microscopy using a FEI Technai 20 transmission electron microscope. The ratio between the diameter of an axon and the mean diameter of the myelinated fiber (G-ratio) was determined on 200 randomly chosen fibers (3-6 animals/ genotype) using Image J software (imagej.net). Age-related ultrastructural changes were identified according to the description provided by Peters and Sethares (Alan Peters and Claire Folger Sethares, The fine structure of the aging brain (www.bu.edu/agingbrain)) and quantified in 15 different microscopic fields /mouse (3-6 animals/genotype; average neurons/genotype 1118; range 929-1498).

#### Immunofluorescence Staining of Brain Sections

Brain sections (30 μm thick) were obtained as described (Chitu et al., 2015) and stained using antibodies to ionized calcium binding adaptor molecule 1 (Iba1) (rabbit IgG; Wako Chemicals, Richmond, VA or goat IgG; AbCam, Cambridge, MA), cystatin F (rabbit IgG; Fisher, Pittsburgh, PA), poly(ADP-ribose) (mouse monoclonal, Millipore, Billerica, MA), myelin basic protein (mouse monoclonal, BioLegend, Dedham, MA), PDGFRα (goat polyclonal, Minneapolis, MN), CC1 (a mouse antibody to APC reacting with Quaking 7, Millipore, Danvers, MA) and NeuN (mouse monoclonal, Millipore, Danvers, MA). Rat anti-P2ry12 was a gift from Dr. Oleg Butovsky (Harvard Medical School). Secondary antibodies, conjugated to either Alexa 488, Alexa 594 or Alexa 647, were from Life Technologies (Grand Island, NY). Images were captured using an Nikon Eclipse TE300 fluorescence microscope with NISElements D4.10.01 software. Quantification of cell numbers was performed manually. Quantification of fluorescent areas was performed using ImageJ. Images were cropped and adjusted for brightness, contrast and color balance using Adobe Photoshop CS4.

#### Microglia morphometry

Morphometric analysis of microglia was carried out on maxiumum intensity projections of Iba-1 stained tissue sections from 3 mice/genotype using FIJI as described (Young and Morrison, 2018). Images were obtained using a Leica SP5 Confocal microscope.

### QUANTIFICATION AND STATISTICAL ANALYSIS

#### Statistical Analyses

Statistical analysis was performed using the GraphPad Prism 7 software (GraphPad, La Jolla, CA, USA). Data were screened for the presence of outliers using the ROUT method, assessed for Gaussian distribution by D’Agostino-Pearson omnibus normality test and analyzed using Student’s t test, one-way ANOVA, the Kruskal-Wallis test or two-way ANOVA, as appropriate. Pairwise differences were identified using post-hoc multiple comparison tests. The level of significance was set at p< 0.05. Data within each group are presented as averages ± standard error of the mean (S.E.M.). Only those differences that have reached statistical significance are indicated on the figures.

### DATA AND CODE AVAILABILITY

#### Data Availability

All data are available in the main text or the supplemental materials. RNA Seq data will be deposited in the Gene Expression Omnibus (GEO) database; accession number pending.

## Supplemental Information

### Supplemental Figures 1-5

**Figure S1.**
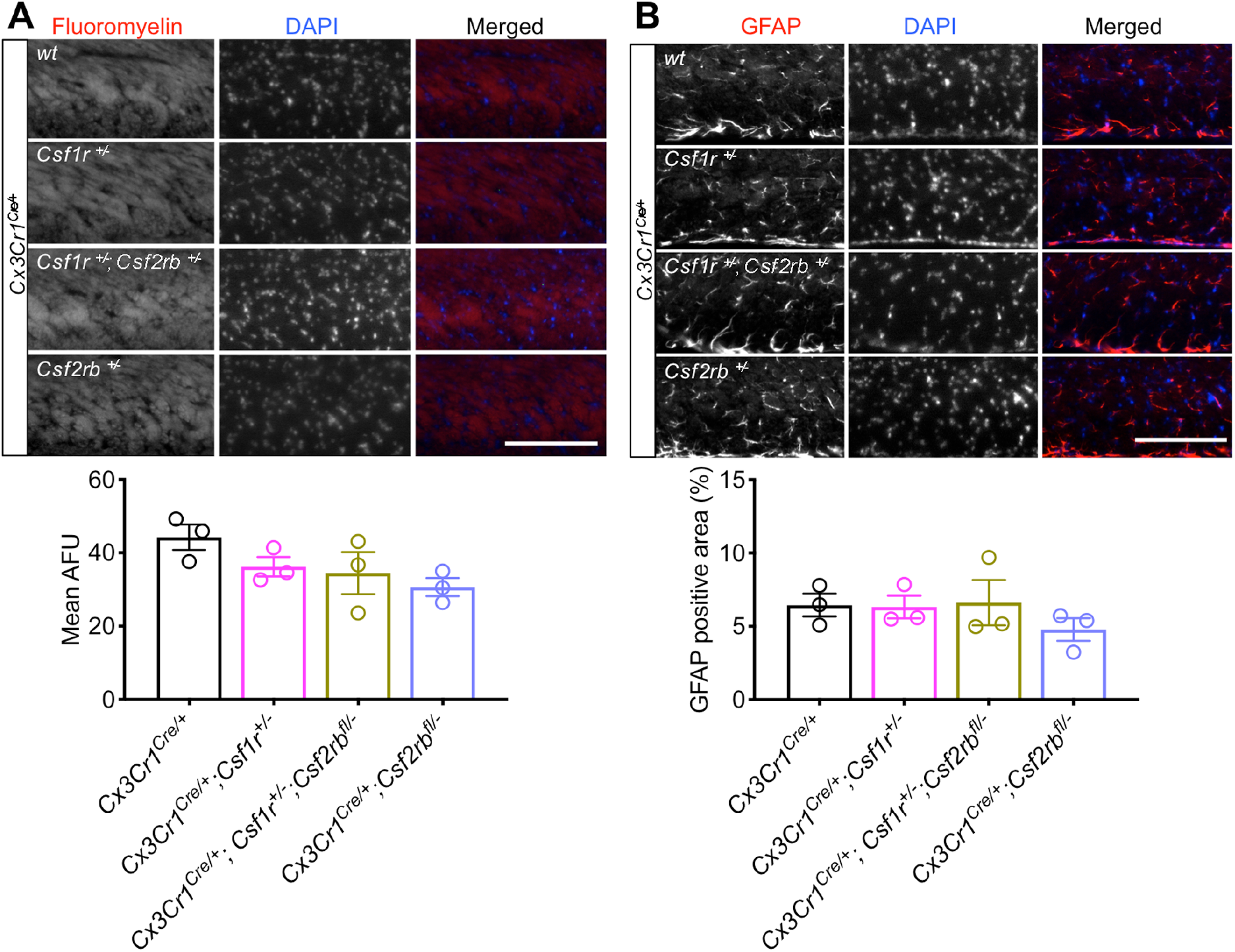
Absence of histopathological changes in the callosal white matter of 3-month-old *Cx3Cr1^Cre/+^; Csflr^+/-^; Csf2rb^fl/+^* mice. (A) Fluoromyelin staining shows normal myelination. (B) Absence of astrocytosis shown by GFAP staining. AFU, arbitrary fluorescence units. Scale bars, 100μm. Data ±SEM; one-way ANOVA not significant. Related to Figure 1.

**Figure S2.**
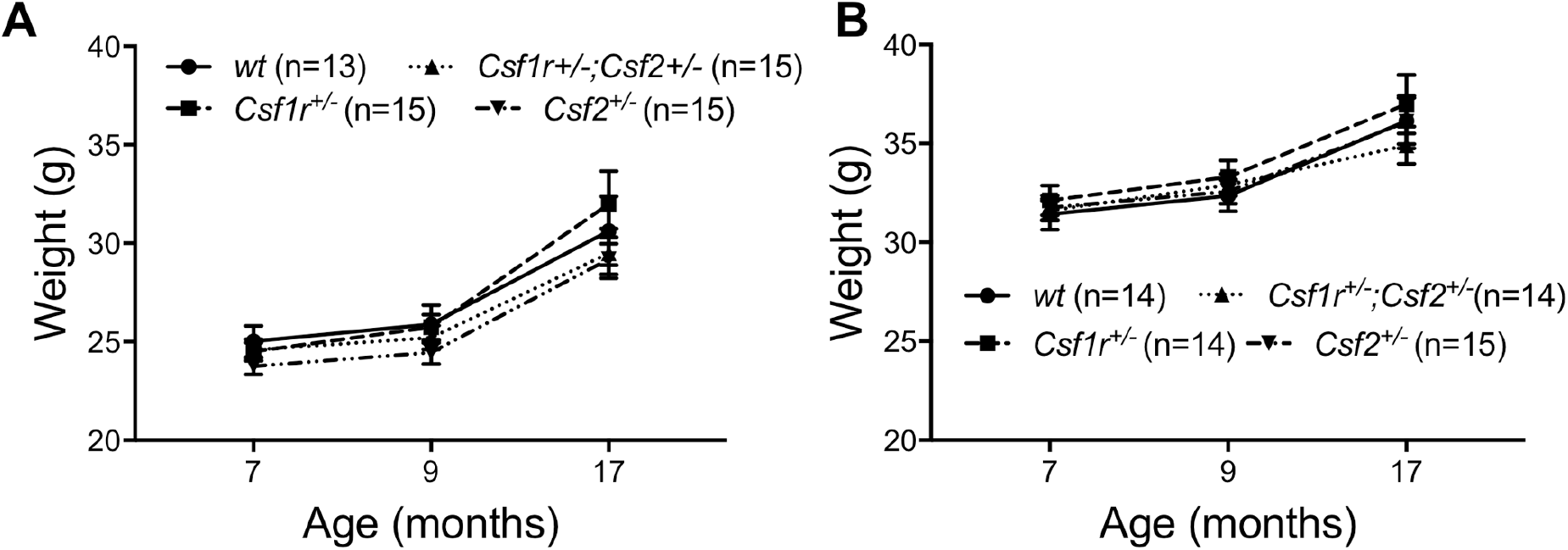
Body weight increases with age. Related to Figures 2 and 3. (A) Female mice. [Genotype x age interaction, F(6, 108) = 1.018; p = 0.4174. Genotype, F(3, 54) = 0.049; p = 0.8028. Age, F(2, 108) = 113.3; p < 0.0001]. (B) Male mice [Genotype x age interaction, F(6, 106) = 0.7987; p = 0.5730. Genotype, F(3, 53)=0.2837; p = 0.8369. Age, F(2, 106) = 113.3; p < 0.0001].

**Figure S3.**
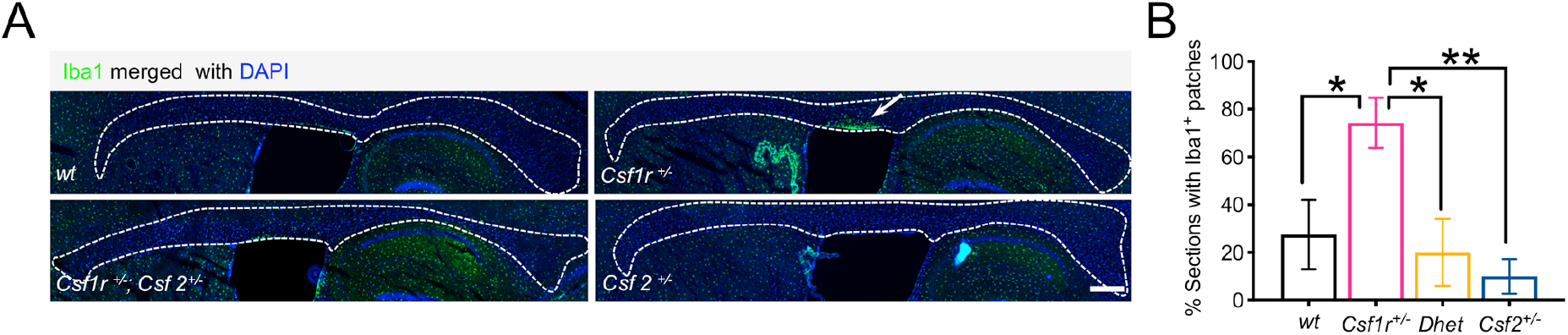
*Csf2* heterozygosity reduces the accumulation of activated microglia in the white matter of *Csf1r^+/-^* mice. Related to Figure 4. (A) A patch of high microglial density in the callosal white matter (dotted lines), located proximal to the lateral ventricle (arrows). Scale bars, 350 μm. (B) Frequency of sections containing microglial patches in wild type, single and double heterozygous mice (percent calculated from 8 sections/mouse, 5 mice/genotype). Means ± S.E.M., one-way ANOVA, p = 0.0075, followed by the Holm-Sidak post-hoc test, *, p<0.05.

**Figure S4.**
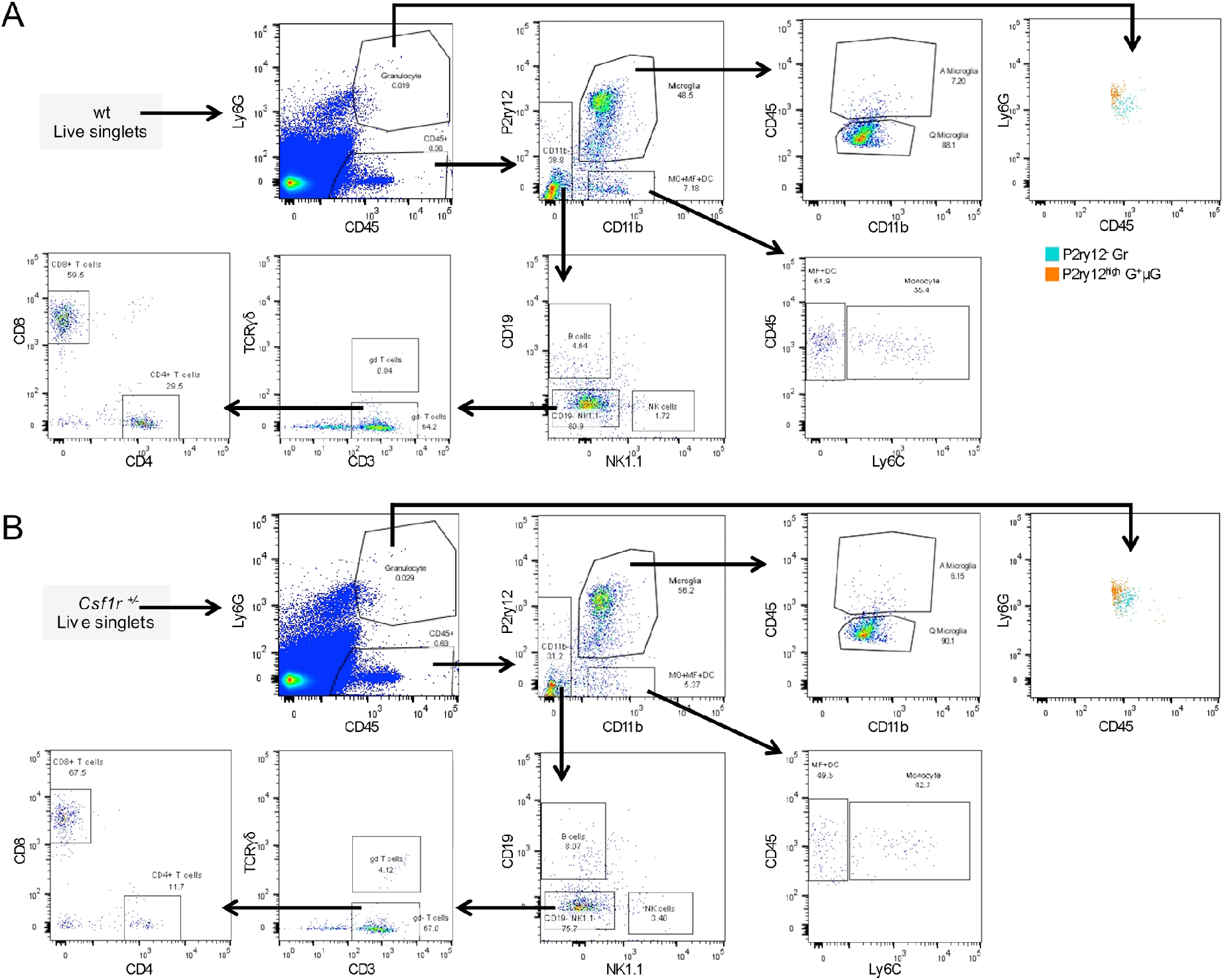
Gating strategy utilized to examine immune infiltrates in the brain by flow cytometry. Related to Figure 4. Panel A shows representative results for wt mice, panel B for Csf1r^+/-^ mice. Because of its heterogeneous nature, the Ly6G^+^ population was further examined for expression of P2ry12 and CD11b. The right panels show that this population contains both P2ry12^-^ (bona-fide granulocytes) and P2ry12^high^ cells, presumably activated microglia.

**Figure S5.**
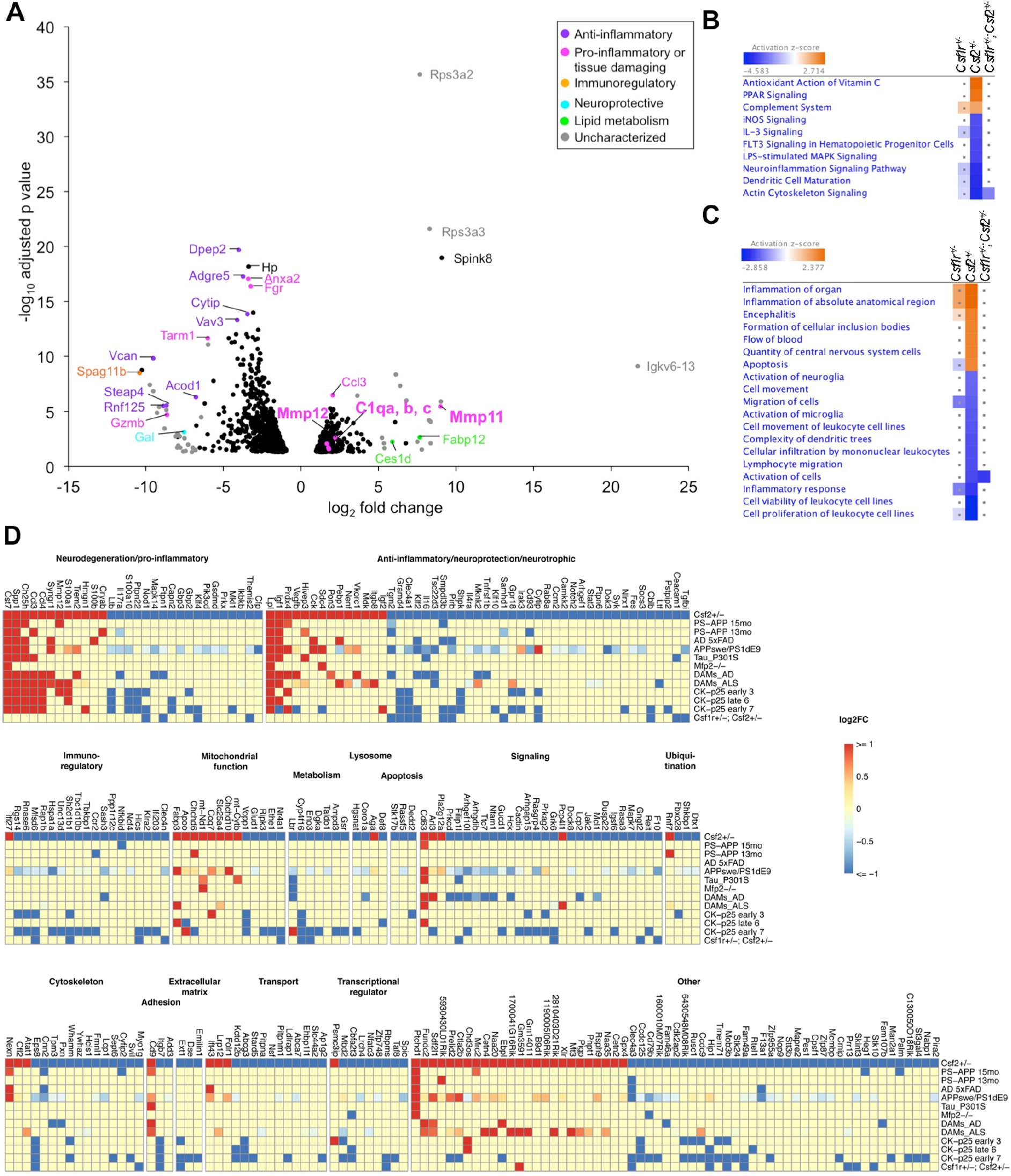
Changes in the microglial transcriptome associated with *Csf2* heterozygosity. (A) Volcano plot highlighting DEGs of interest in *Csf2^+/-^* microglia. (B and C) IPA-generated list of pathways (B) and biological processes (C) uniquely affected by *Csf1r* heterozygosity and their predicted activation status in *Csf1r^+/-^* and double heterozygous microglia. Dots indicate no significant difference. (D) Heatmap showing the overlap of *Csf2^+/-^* DEGs with genes differentially expressed in other mouse models of neurodegenerative disease.

**Supplemental Table 1.**
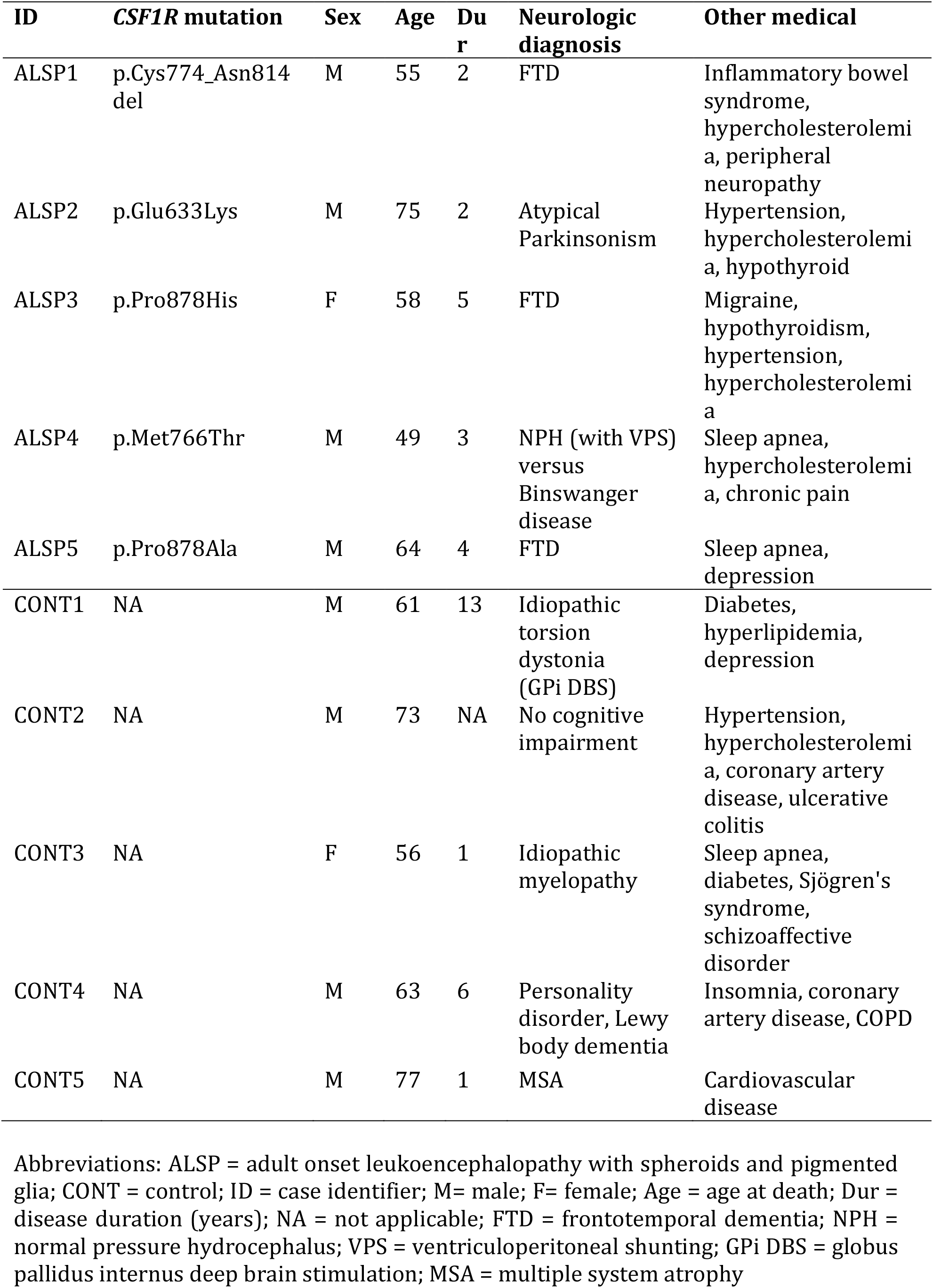
Summary of Subjects Analyzed for *CSF2, CST7* and *CH25H* Gene Expression. Related to Figures 1 and 6.

| ID | *CSF1R* mutation | Sex | Age | Dur | Neurologic diagnosis | Other medical |
| --- | --- | --- | --- | --- | --- | --- |
| ALSP1 | p.Cys774_Asn814 del | M | 55 | 2 | FTD | Inflammatory bowel syndrome, hypercholesterolemia, peripheral neuropathy |
| ALSP2 | p.Glu633Lys | M | 75 | 2 | Atypical Parkinsonism | Hypertension, hypercholesterolemia, hypothyroid |
| ALSP3 | p.Pro878His | F | 58 | 5 | FTD | Migraine, hypothyroidism, hypertension, hypercholesterolemia |
| ALSP4 | p.Met766Thr | M | 49 | 3 | NPH (with VPS) versus Binswanger disease | Sleep apnea, hypercholesterolemia, chronic pain |
| ALSP5 | p.Pro878Ala | M | 64 | 4 | FTD | Sleep apnea, depression |
| CONT1 | NA | M | 61 | 13 | Idiopathic torsion dystonia (GPi DBS) | Diabetes, hyperlipidemia, depression |
| CONT2 | NA | M | 73 | NA | No cognitive impairment | Hypertension, hypercholesterolemia, coronary artery disease, ulcerative colitis |
| CONT3 | NA | F | 56 | 1 | Idiopathic myelopathy | Sleep apnea, diabetes, Sjögren's syndrome, schizoaffective disorder |
| CONT4 | NA | M | 63 | 6 | Personality disorder, Lewy body dementia | Insomnia, coronary artery disease, COPD |
| CONT5 | NA | M | 77 | 1 | MSA | Cardiovascular disease |

Abbreviations: ALSP = adult onset leukoencephalopathy with spheroids and pigmented glia; CONT = control; ID = case identifier; M= male; F= female; Age = age at death; Dur = disease duration (years); NA = not applicable; FTD = frontotemporal dementia; NPH = normal pressure hydrocephalus; VPS = ventriculoperitoneal shunting; GPi DBS = globus pallidus internus deep brain stimulation; MSA = multiple system atrophy

## Supplemental tables not included in the main PDF

Supplemental Table 2: Transcriptomic changes in Csf1r+/-, Csf2+/- and Csf1r+/-; Csf2+/- microglia relative to wt. Related to Figure 5 and Supplemental Fig. S5.

Supplemental Table 3: Gene ontology clustering. Part 1: Gene ontology clustering of transcripts upregulated in Csf1r+/- microglia. Part 2: Gene ontology clustering of transcripts downregulated in Csf1r+/- microglia. Related to Figure 5.

Supplemental Table 4: Relative expression of neurotrophic factor transcripts in Csf1r+/-, Csf2+/- and Csf1r+/-; Csf2+/- microglia. Related to Figure 5.

Supplemental Table 5: Canonical pathways affected by Csf1r or Csf2 heterozygosity. Part 1: Canonical pathways affected by Csf1r in microglia and effect of Csf2 or combined (Dhet) heterozygosity on the same pathways. Part2: Canonical pathways specifically affected by Csf2 heterozygosity in microglia and effect of combined (Dhet) heterozygosity on these pathways.

Supplemental Table 6: Biological processes affected by Csf1r or Csf2 heterozygosity. Part1: Biological processes affected by Csf1r heterozygosity in microglia and effect of Csf2 or combined (Dhet) heterozygosity on the same processes. Part 2: Biological processes specifically affected by Csf2 heterozygosity in microglia and effect of combined (Dhet) heterozygosity on these processes. Related to Figure 5.

Supplemental Table 7. Genes co-regulated in Csf1r+/- microglia and other models of neurodegenerative disease. Related to Figure 5.

Supplemental Table 8. Genes co-regulated in Csf2+/- microglia and other models of neurodegenerative disease. Related to Figure S5.

## Abbreviations

CSF-1 R: Colony stimulating factor-1 receptor
CSF-2: Colony stimulating factor-2
ALSP: Adult-onset leukoencephalopathy with axonal spheroids and pigmented glia

